# A graph-based practice of evaluating collective identities of cell clusters

**DOI:** 10.1101/2024.06.28.601289

**Authors:** Yuji Okano, Yoshitaka Kase, Hideyuki Okano

## Abstract

The rise of single-cell RNA-sequencing (scRNA-seq) and evolved computational algorithms have significantly advanced biomedical science by revealing and visualizing the multifaceted and diverse nature of single cells. These technical advancements have also highlighted the pivotal role of cell clusters as representations of biologically universal entities such as cell types and cell states. However, to some extent, these clusterings remain dataset-specific and method-dependent. To improve comparability across different datasets or compositions, we previously introduced a graph-based representation of cell collections that captures the statistical dependencies of their characteristic genes.

While our earlier work focused on theoretical insights, it was not sufficiently adapted and fine-tuned for practical implementation. To address this, the present paper introduces an improved practice to define and evaluate cellular identities based on our theory. First, we provide a concise summary of our previous theory and workflow. Then, point-by-point, we highlight the issues that needed fixing and propose solutions. The framework’s utility was enhanced by leveraging alternative formats of cellular features such as gene ontology (GO) terms and effectively handling dropouts. Supplemental techniques are offered to reinforce the versatility and robustness of our method.

## Introduction

It has been more than 10 years since the birth of scRNA-seq [1], and the technology is now recognized as a prominent game-changer in molecular biology. Like the pioneering technologies in this field—DNA micro array and bulk RNA-seq—scRNA-seq can observe multidimensional gene expression profiles. Additionally, it can provide such information at the single-cell-level. The abundance of information it provides has contributed to revealing the detailed biology of various cell types, but the excessive resolution has blurred the conceptual boundary between static cell types and transient cell statuses [2]. Consequently, “clusters”—chunks of samples sharing similar geometrical properties in the data space—have overwritten the classical notion of cell types. The formation of cell clusters depends on the sampling stochasticity of the dataset, and their biological properties might deviate from the original doctrine of cell types [3]. Therefore, a theoretical backbone and an effective method to link theory and experimental data are essential to distinguish universal characteristics of specific samples from piles of extrinsic noise.

In our previous research, we proposed a gene regulatory network (GRN)-based representation of cell clusters, in which the edges of GRNs explain statistical dependencies between two genes. This demonstrated that the similarity of two clusters can be defined as a quasi-pseudo-metric function *d*^*^ [3]. To discuss the mathematical properties of the quasi-pseudo-metric space of cell clusters, we defined the novel terms “eigen-cascades” and “cell class,” and introduced their algebraic structures. Eigen-cascades refer to a set of marker genes and pairs of genes that are statistically interdependent (i.e., isomorphic to the direct sum of the vertex set and the edge set of a GRN). A cell class refers to a cell cluster characterized by the corresponding eigen-cascades. Note that the nuances of cell cluster and cell class definitions are slightly different, even though we might use those terms interchangeably in this article (See Appendices for more explanation). When two cell classes ∀[*x*], [*y*] are represented by the GRNs regarding a set of genes *G*, and the two GRNs (eigen-cascades) are denoted as *C*_[*x*]_(*G*) and *C*_[*y*]_(*G*), a bivariate function *d*^*^ that maps a pair of cell classes to real numbers is defined as follows [3]:

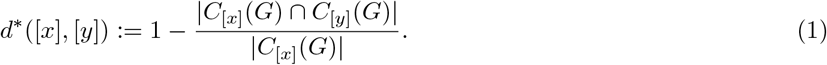

Eq. (1) is derived from the Hamming distance function (a metric function that measures the difference between two character strings) and modified to embrace the tendency of the Peter and Clark (PC) algorithm, one of the simplest Bayesian network algorithms [3, 4, 5]. With these concepts, we also proposed frameworks to compare the similarities of two given cell clusters. Our scheme comprises two fundamental steps: (i) formation of GRNs and (ii) evaluation of their similarity (Figure 1A). As the definition of cell classes is independent of the choice of data analysis methods, this framework can be applied into various cases regardless of any feature engineering (such as the data preprocessing) or clustering methods.

**Figure 1:**
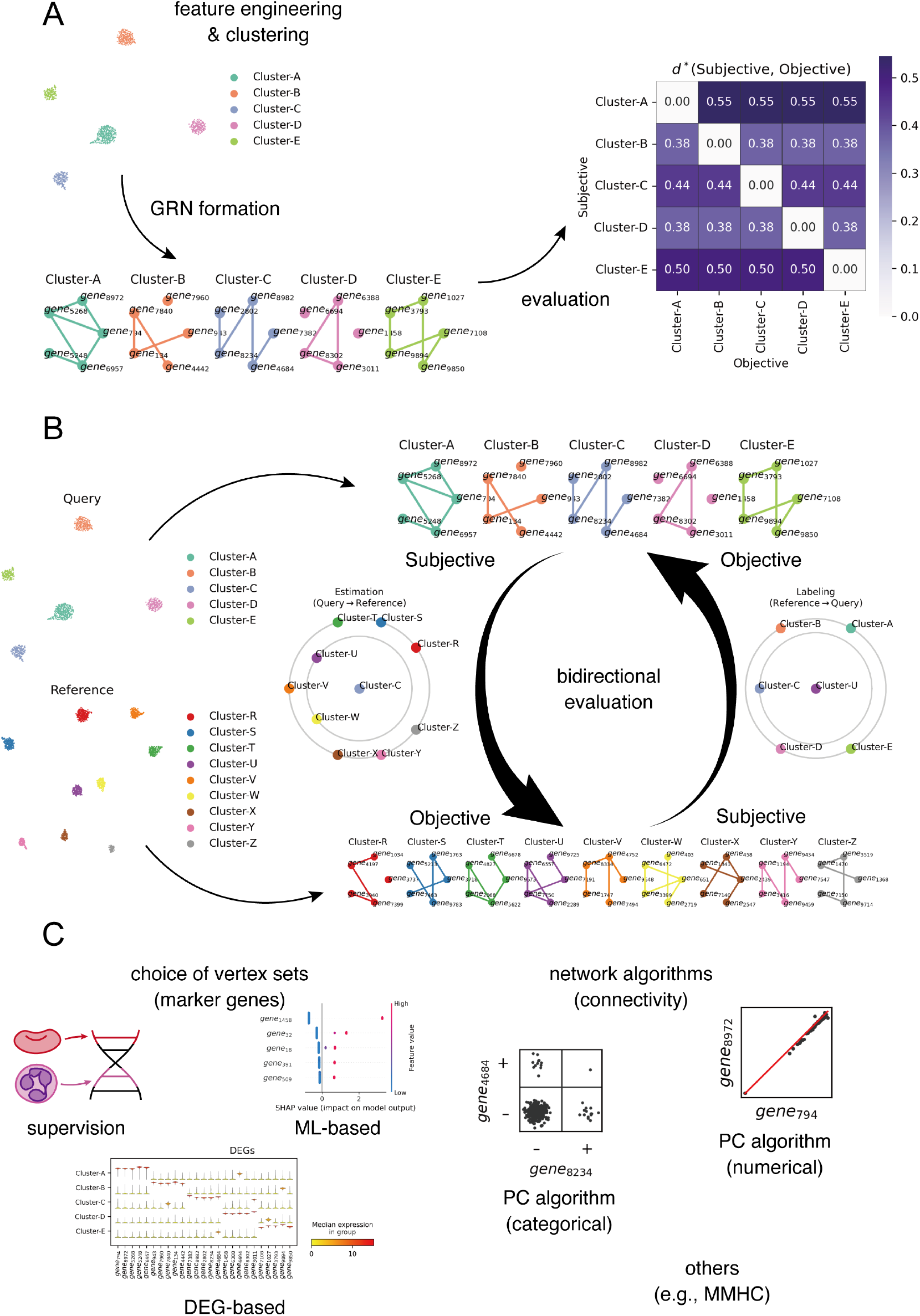
The framework of the GRN-based characterization and annotation of cell classes. **A**: The foundation of the GRN-based characterization of cell classes. After clustering in the designated data space by arbitrary methods, cell classes (the clusters) can be represented by GRNs of corresponding genes of choice. The similarity of two GRNs with the same vertex (marker gene) is evaluated with the asymmetrical function *d*^*^, where the return values reflect the similarity from the viewpoint of the subjective clusters. **B**: Schematic of the GRN-based scRNA-seq data annotation. Expecting the referential data to reflect canonical states of target sample domains, the evaluation of the similarity among cell classes can be performed bidirectionally. **C**: Methodological variations of the selection of vertex sets (marker genes) and the algorithms to compute the network structures of GRNs.

The framework can be applied into the annotation of scRNA-seq data when a referential dataset is available (Figure 1B). Because annotation is an act of tagging clusters with descriptions in natural languages, the biological features of annotated clusters are often treated as common and preserved properties of the cell types after which the clusters are named. Accordingly, it is better to have a large enough referential dataset reflecting canonical states of specific cell types shared with the query dataset. Using GRN-based characterization, cell classes are annotated with the name of the most similar cell class; however, comparisons of cellular identities can be bidirectional due to the asymmetry of *d*^*^. We refer to the similarity of cell classes from the perspectives of the query data as “estimation,” and the one from the point of view of the referential data as “labeling.” These GRN-based annotations can be visualized with planet plots, in which the subjective cell class (denote here as [*x*]) is located in the center and the radii of the circles reflect *d*^*^([*x*], ·) values for all cell classes placed on the circumferences.

The performance of those frameworks built around GRNs has a bottleneck in the GRN creation step, which involves configuring the vertex sets and choosing of the network algorithm (Figure 1C). Similar to feature engineering and clustering methods, each step of GRN formation offers various options. In our last paper, we introduced a combined method of manual curation using review articles and a machine learning (ML)-based feature selection with a gradient boosting decision tree (GBDT) model incorporating L1 and the L2 regularizations [3]. For manual supervision, GO terms can serve as an additional information source. However, prior identification of sample components is essential to create meaningful GRNs by integrating domain-specific information. The differentially expressed gene (DEG)-based method can be a more heuristic and a less interactive option since the differential expression analysis (DEA) semi-automatically selects DEGs. Regarding the network algorithms, there are several possible options. In our previous paper, we implemented our codes using the numerical (i.e., correlation-based) PC algorithms provided in Pgmpy [6], a Python package for probabilistic graphical models. Another variation of the PC algorithm based on the chi-square test, suitable for categorical data, is also a realistic option when the expression values can be binarized. We also mentioned that the max-min hill-climbing (MMHC) algorithm, which combines constraint-based and scoring-based methods [7], is one of the promissing alternatives to the PC algorithm.

So far, we have highlighted the versatility of our framework by providing examples that demonstrate its applicability across various data analysis methods. Our intention was to allow researchers to integrate their expertise in specific sample domains, or preferences, by textually describing the samples in biological terms. This customization ensures that the metrics for cellular identities are crafted to align with the specific research scopes, providing both necessary and sufficient resolutions. However, this design choice has the drawback of making our algorithm less user-friendly, as it requires a significant amount of effort in annotation, even when annotation might not be a primary focus of their projects. To validate our theory across a wide range of cases, it is essential to refine the practices related to GRN-based annotation and streamline the overall workflow.

In this article, we address three major issues where the former protocol left room for improvement and provide more practical solutions for each while leveraging the backbone theory of GRN-based comparisons of cluster-wise cellular identities (i.e., cell classes).

## Results

### Challenges for the framework of GRN-based methods

To clarify our objective for this article, this section describes the three major challenges for the practical use of GRN-based annotation:

#### 1. Difficulty of effective gene selection

In our previous study, we proposed a method for selecting marker genes that combines supervised curation, based on a review paper, with ML-based feature selection, leveraging the feature importance in an L1-regularized GBDT model. While we mentioned that various options exist for marker gene selection, each strategy has its unique drawback.

Supervision by the experimenter struggles with completeness and arbitrariness, even when citing reliable sources (e.g., review papers or GO terms). For example, in our previous paper, we selected *SLC1A2, VIM*, and *AQP4* as glial markers based on a review article [8]. These glial markers are associated with various GO terms, but other genes are also tagged with these terms (Figure 2A). This highlights the incompleteness of these three genes in representing all aspects of glia. Furthermore, the many-to-many correspondence between genes and GO terms makes it challenging to draw a clear line between adopted marker genes and others.

**Figure 2:**
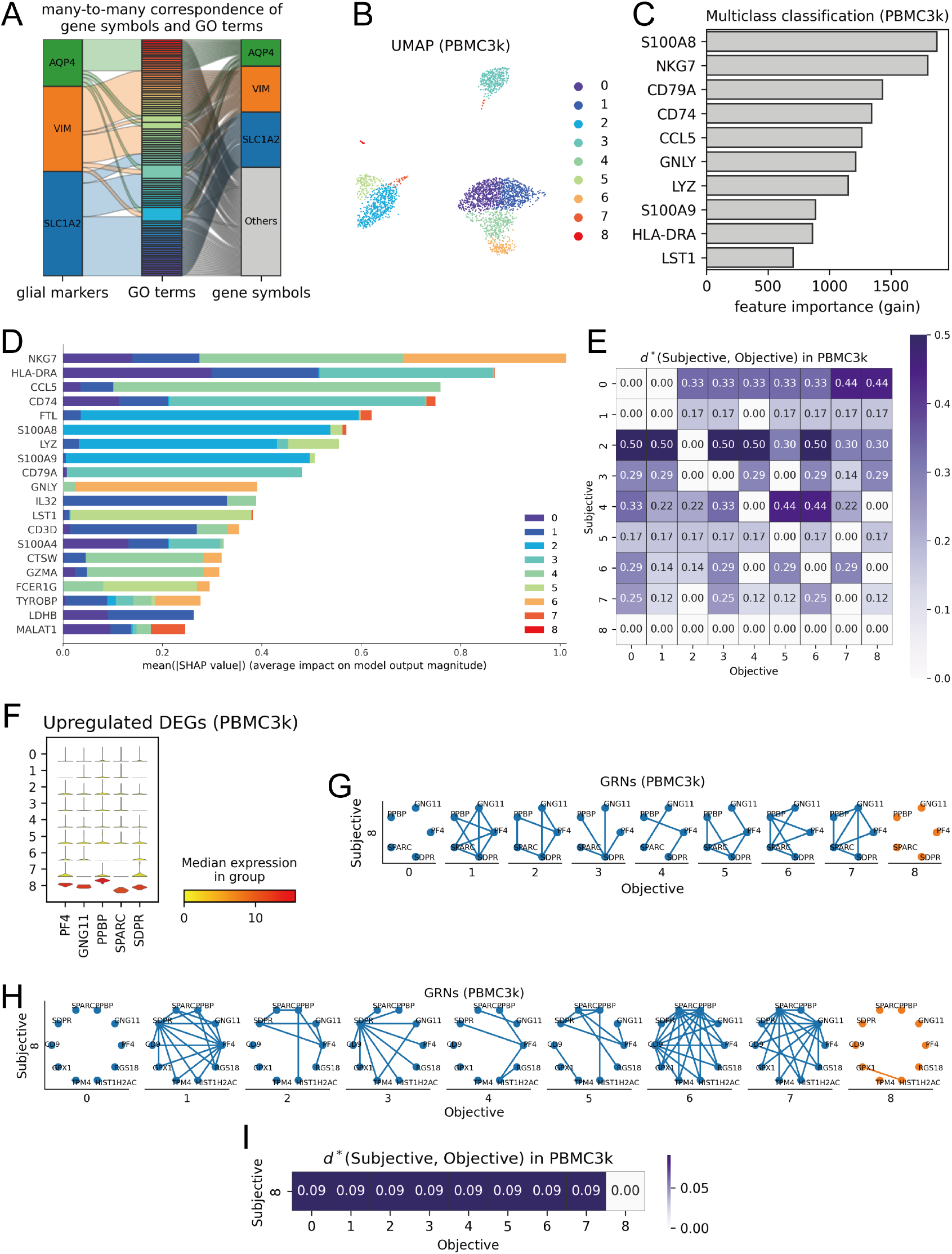
Examples of the major issues on the GRN-based frameworks. **A**: Alluvial plot showing the many-to-many correspondence between gene symbols and GO terms. **B**: UMAP of the PBMC3k dataset. The markers are colored according to the cluster. **C**: The top 10 genes with the highest feature importance according to the multiclass classification LightGBM model. **D**: The top 20 genes with the highest mean SHAP values according to the multiclass classification LightGBM model. **E**: The *d*^*^ values based on the GRNs of the top 5 DEGs. The rows correspond to the subjective cell classes, and the columns correspond to the objective ones. **F**: The top 5 DEGs of cluster 8. **G**: The GRNs of the clusters 0∼8 based on the top 5 DEGs of cluster 8. **H**: The GRNs of the clusters 0∼8 based on the top 10 DEGs of cluster 8. **I**: The re-calculated *d*^*^ values among the GRNs based on the top 10 DEGs of cluster 8.

The ML-based approach is another method we implemented in our last report, and it also has its unique problems. As shown in Figure S1, the standard workflow of scRNA-seq data analysis involves several steps: data quality control (QC); normalization, including reads per million (RPM) transformation and logarithmic transformation; highly variable gene (HVG) extraction; dimensionality reduction, such as principal component analysis (PCA), truncated singular value decomposition (TSVD), uniform manifold approximation and projection (UMAP) [9], etc.; clustering; DEA; annotation; and other downstream analyses [10]. Creating an effective ML model requires considerable time beyond actual run times for fine-tuning model configurations, which can be excessively effortful for gene selection for GRNs, especially when annotation is not the primary goal of the data analysis. Moreover, even with a well-performing ML model, extracting informative features can suffer from arbitrariness in the selection. To illustrate these difficulties, we analyzed an open-source scRNA-seq dataset of peripheral blood mononuclear cells (referred to as “PBMC3k” for convenience), distributed bt the company 10X Genomics. Starting from QC, we proceeded to leiden clustering, resulting in 9 clusters (0∼8) as shown in Figure 2B. We then created a GBDT model for multiclass classification, predicting clusters from gene expression values. The model performed well in terms of the area under the curve (AUC) of the one-versus-rest (OvR) receiver operating characteristic (ROC) curves; the macro average of the OvR ROC curves; the average precision (AP) of the OvR precision-recall (PR) curves; the micro average of the OvR PR curves; and the accuracy score (Figure S2A-D). In our previous article, we created a three-class classification model and used feature importance as a criterion (Figure 2C). However, this approach did not work for the nine-class classification model due to the lack of clear boundaries between key and negligible features, even when selecting the top 10 features of importance. Since GRNs require pairwise edge calculation, modelers should avoid using an excessive number of genes for computational efficiency. Besides feature importance scores, Shapley additive explanations (SHAP) scores can be an alternative metric to visualize the correspondence between features and classifications [11]. Despite SHAP scores providing more intuitive and precise explanations (Figure 2D and S3A-I), it remains challenging to introduce objective thresholds for gene selection due to the drastically varying distributions of SHAP scores across different classes. Consequently, the ML-based approach is not the most effective way to select marker genes for representing cell classes, as it requires subjective and case-by-case decisions on the number of marker genes to adopt, making it more time-consuming. ML-based approaches might work well if the character of the samples is completely unknown or the consensus among experts is yet to be settled. However, even under such conditions, alternative methods such as the DEG-based approach should be considered.

The DEG-based approach is another alternative that can be smoothly integrated into the regular scRNA-seq data analysis pipeline. Despite its heuristic nature and promptness, this method also has its shortcomings. Using the PBMC3k data processed the same way as in the previous section, we will illustrate this with an example. The GRN-based approach offers the advantage of a swift procedure by directly applying the top DEGs into the vertex sets. Accordingly, we applied the top 5 DEGs of each cluster to the vertex sets (Figure S4A), created GRNs based on those genes (Figure S4B), and calculated the *d*^*^ values (Figure 2E). Observing the bottom row of the heatmap, the *d*^*^ values were all zero from the perspective of cluster 8 even though it showed significantly different expression patterns of the top 5 marker genes (Figure 2F). Additionally, the top 30 upregulated GO terms of each cluster indicate that cluster 8 could exclusively be annotated as “megakaryocytes,” while the others exhibited different cellular characters (Figure S5A-I). This indicated that the GRNs did not work properly for identifying cluster 8, as the zero *d*^*^ values for the clusters 0∼7 implied that these clusters and cluster 8 were indistinguishable in terms of the GRNs with the given vertex sets. Increasing the number of DEGs to 10 added an edge to the GRN for cluster 8, resulting in non-zero *d*^*^ values (Figure 2H-I). As the GO terms suggested no other clusters of megakaryocytes except cluster 8, the new *d*^*^ values appeared to correctly indicate that the clusters 0∼7 were equally different from cluster 8. Thus, the marker-gene selection is an intricate step that requires repetitive adjustments and validations especially when exploring the optimal number of the DEGs to use for the vertex sets.

In this paragraph, we have highlighted issues with various marker-gene selection methods and emphasized that selecting necessary and sufficient genes to represent certain cellular identities is time-consuming and sometimes computationally intensive, though it is a key part of GRN-based annotation. Given this, and considering that annotation is often not the ultimate goal of the scRNA-seq data analyses, developing an alternative method for finding marker genes is necessary to reduce computational costs and streamline the time required before initiating the main analyses.

#### 2. Statistical issue: independence v.s. uncorrelation

The statistical independence of two events *A* and *B* is defined as a situation where the following equation holds:

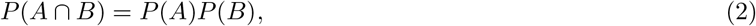

where *P* (·) is the probability of an event. On the other hand, the correlation coefficient *Corr*(*X, Y*) of stochastic variables *X* and *Y* is defined as follows:

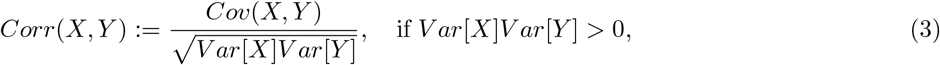

where *E*[·] is the expected value, *V ar*[·] is the variance, and *Cov*(·, ·) is the covariance. Independent variables exhibit a correlation coefficient of zero, but the converse is false. For example, if *X* ∼ *U* (−1, 1), where *U* (−1, 1) refers to the uniform distribution over the interval from −1 to 1, then *Corr*(*X, X*^2^) = 0 although *X* and *X*^2^ are dependent. Therefore, strictly speaking, it is not appropriate to substitute the chi-square test or the exact test with the t-test of correlation. Furthermore, the correlation-based method does not work well when the gene expression matrices are highly sparse regarding the selected genes. As Eq. (3) holds if, and only if, both *V ar*[*X*] and *V ar*[*Y*] are non-zero values, under circumstances where all samples in a cluster exhibit zero counts for certain genes required in the vertex set, the correlation-based approach is inappropriate. This situation is by no means an unrealistic hypothetical counterexample. For example, a phenomenon called “dropout” is a characteristic of scRNA-seq data where gene expressions are not detected due to the inefficiency and the stochasticity of scRNA-seq [12], resulting in high sparsity of scRNA-seq data matrices.

In our previous work, we introduced a correlation-based algorithm to construct GRNs, compromising rigor to accommodate continuous gene expression values. To address this issue in the current study, we propose an effective method to binarize the gene expression values, allowing the new algorithm to rely on statistical tests of independence. This update will better aligh our algorithm with the original concept of our theory.

#### 3. Insufficient responsiveness to gene expression values

GRNs have originally been designed to represent cellular functions by establishing edges between two statistically dependent genes. The correlation-based GRN generation follows the same idea, drawing edges between two vertexes where correlations exist. Although these strategies can visualize the co-occurrence or mutual exclusivity of the gene expressions, actual expression values are dismissed. This failure leads to misassignments of cellular identities in practical cases, as demonstrated with the PBMC3k example; namely, the GRNs of clusters 0 and 8 showed identical structures despite significantly different expression patterns of the marker genes forming the vertex set (Figure 2F-G). This example highlights not only the difficulty of the gene selection but also the insufficient responsiveness of GRNs to gene expression values.

### Semi-automated marker-gene suggestion

Manual curation of marker genes for GRNs struggles with arbitrariness and incompleteness, while semi-automated ML-based and DEG-based methods often require overly recurrsive trials to find optimal sets of marker genes. To improve workflow efficiency, we developed an algorithm to automatically suggest similar genes to supplement given marker genes.

We leveraged overlapping GO terms of the given marker genes and mapped them back to gene symbols. For instance, the three glial marker genes in our example share two GO terms in their intersection (Figure 3A), and the similarities of their GO terms can be set-theoretically defined with Jaccard index values (Figure 3B). We interpreted that: 1) the intersection of the Venn diagram contained the pivotal GO terms that reflected the biological semantics collectively defined by the given marker genes; and 2) the minimal Jaccard index value was the indicator of the similarity about the group of genes (therefore it could work as a threshold of acceptance when other genes are added). To find new genes without altering the biological meaning of the list, we querried gene symbols tagged with the pivotal GO terms (Figure 3C), filtered out genes that exhibited lower Jaccard index values compared to any gene in the original list (Figure 3D), and used the remainders to form the new gene list (Figure 3E).

**Figure 3:**
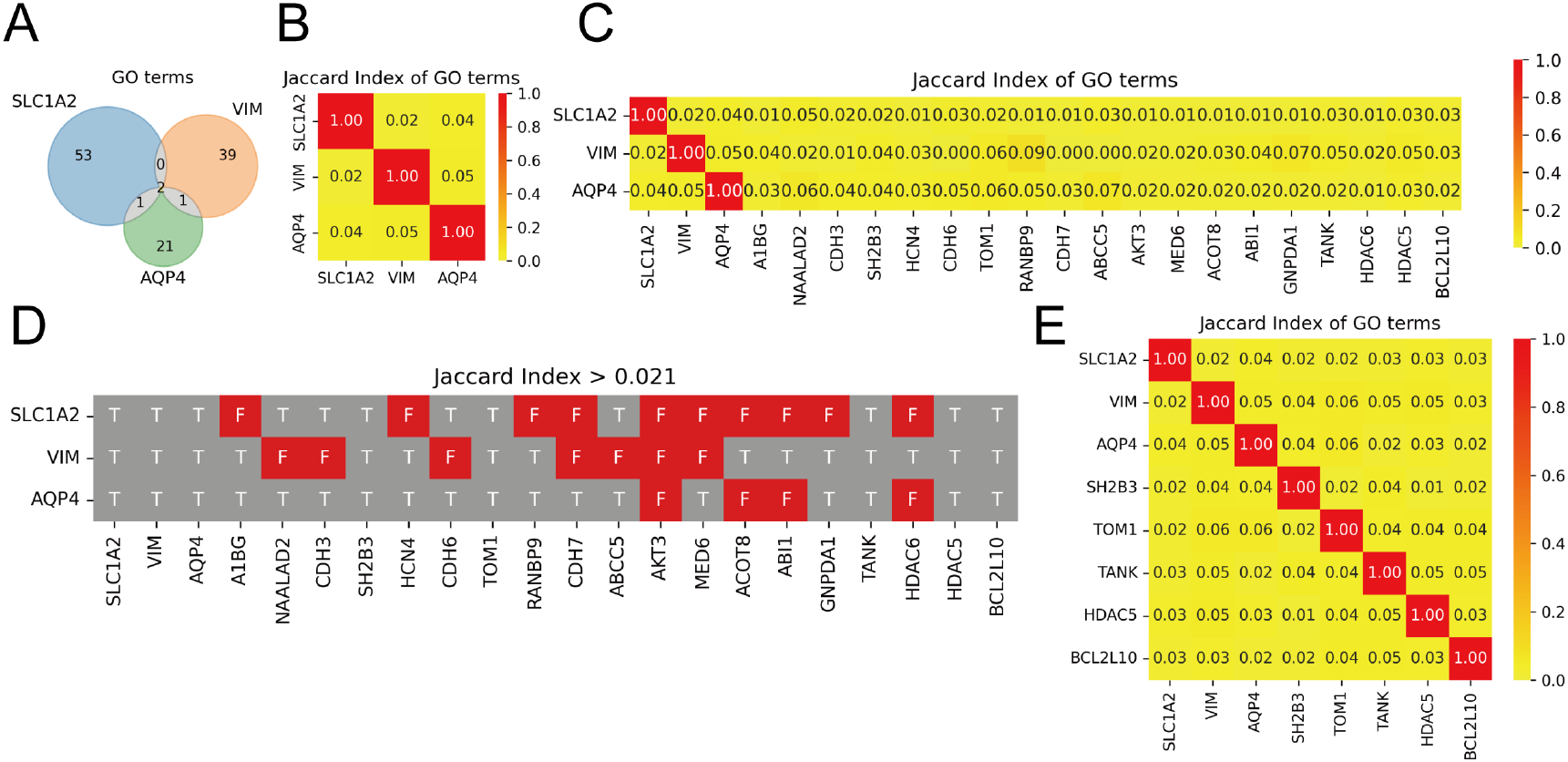
Jaccard index based automated marker gene suggestion. **A**: Venn diagram of the GO terms related to the three glial marker genes. Here, we considered the intersection of the three sets as the pivotal GO terms defined by the three marker genes. **B**: Jaccard index values of the GO terms related to the three glial marker genes. The minimal value was adopted as the threshold for automated gene selection. **C**: Jaccard index values of the GO terms related to the three glial marker genes and other gene symbols subscribed to the pivotal GO terms. **D**: Jaccard index values smaller than the threshold are shown in red, and the others are shown in gray. **E**: Jaccard index values of the GO terms related to the gene symbols included in the output gene list.

We also implemented a combined method of manual and ML-based marker gene selection on the GRN-based annotation using a referential dataset. This labeling evaluates the *d*^*^ values from the referential clusters to the query clusters. Such methods that require manual assignment of marker genes are suitable for characterizing clusters of known cellular identities (i.e., pre-annotated clusters in referential datasets). Accordingly, our new proposal can be applied to similar cases.

### Dropout-based binarization

As discussed, the risk of dropout highlights the pitfalls of the correlation-based algorithms. However, recent studies show that the zero inflation is closely related to data attributes like cell types [12, 13]. Building on this, we consider dropouts as potential indicators of enriched gene expressions. Our binarization algorithm marks non-zero expression values as positive and zeros as negative. For instance, the top two DEGs of cluster 8 in PBMC3k, *PF4* and *GNG11* (both megakaryocyte markers [14]), were rarely expressed in cluster 5 (Figure 2F). Consequently, the 2 *×* 2 contingency table identified the majority of cluster 5 as double-negative (Figure 4A).

**Figure 4:**
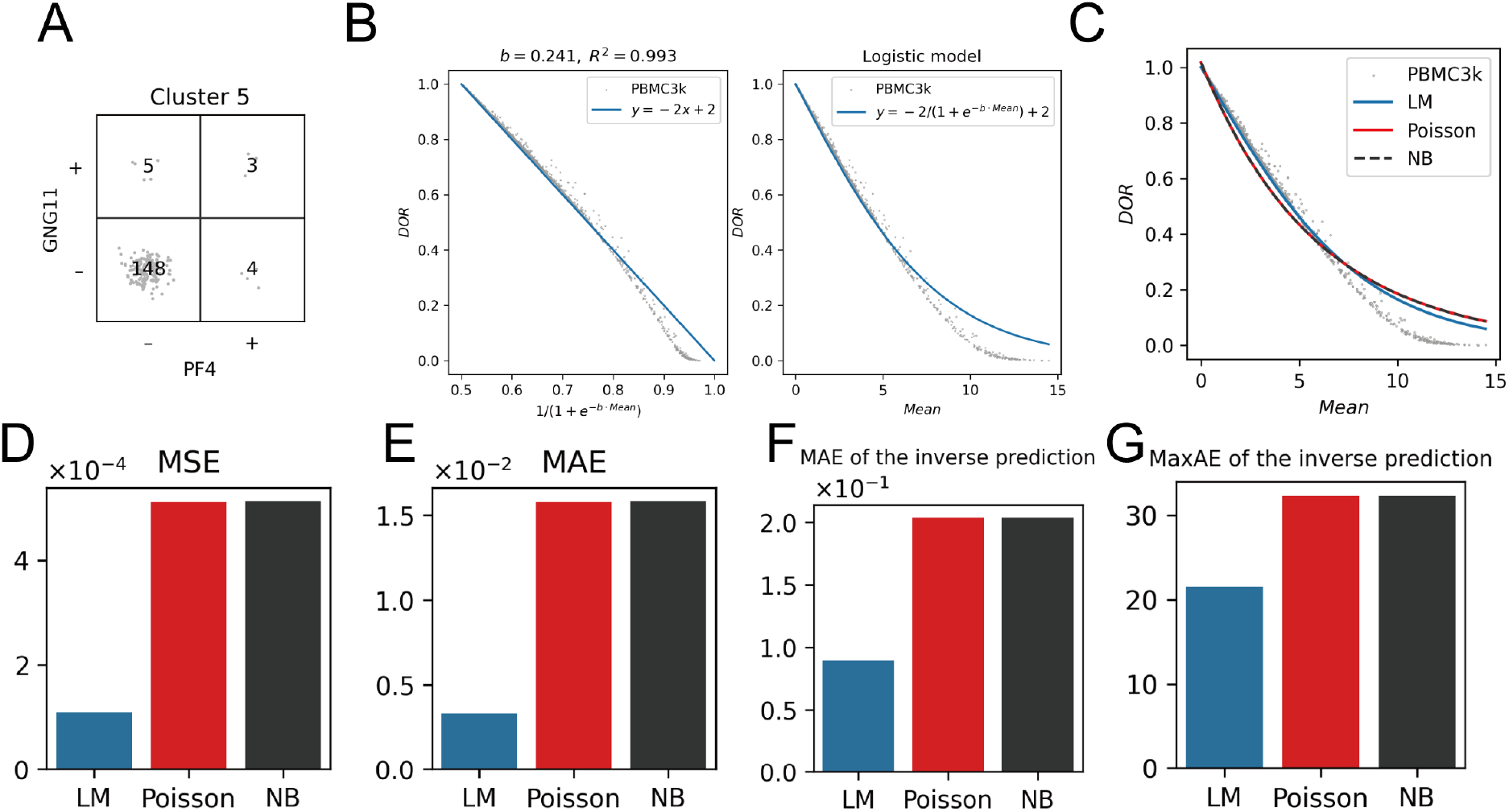
Dropout-based binarization and empirical investigations on DOR. **A**: A dropout-based 2 *×* 2 contingency table of *PF4* and *GNG11* for cluster 5 in PBMC3k (+, non-zero expression values; −, zeros). **B**: The LM of DOR. **C**: The performance comparison with the Poisson regression model (Poisson) and the negative-binomial regression model (NB). **D**: Performance comparison of LM, Poisson, and NB with MSE values. **E**: Performance comparison of LM, Poisson, and NB with MAE values. **F**: Performance comparison of the inverse predictions of LM, Poisson, and NB with MAE values. **G**: Performance comparison of the inverse predictions of LM, Poisson, and NB with MaxAE values.

While we consider dropout a practically useful feature, some experts oppose this and have developed dropout imputation algorithms [15]. To clarify our point, we here examine how dropouts explain the data features.

First, we validated whether the dropout rate (DOR)—the proportion of zeros in the count data of a gene—was associated with the mean values (*log*_2_(*RPM* + 1)) in the PBMC3k data. Although the DOR values and the mean values exhibited a non-linear correspondence, we could successfully establish a linear formulation with a simple logistic transformation on the mean expression values (See Appendices for details). The logistic-transformed mean values fitted well to the linear calibration curve, achieving a coefficient of determination (*R*^2^) of 0.993. Additionally, the inverse-transformed curve aligned well with the data distribution in the scatter plot of the mean values and the DOR (Figure 4B). Hence, we could demonstrate that the DOR values are closely related to the mean expression values, which are the most frequently used summary statistics. As the DOR values are comparable across different datasets, while the mean expression values are unsuitable for trans-dataset comparison, it was suggested that the potential of the calibration curve of DOR and the mean expression could work effectively in cell class comparison using multiple data sources by interchangeably translating the comparable feature and the non-comparable but meaningful one. To benchmark the performance of the model, we coined the name “logistic model (LM)”, and compared it with a Poisson regression model and a negative binomial (NB) regression model (Figure 4C), which are well-known models for explaining dropout events [16]. The mean squared error (MSE) scores of those models indicated that the LM best fitted to the PBMC3k dataset compared to the other competitors (Figure 4D), and its mean absolute error (MAE) (i.e., expected prediction error) in DOR value turned out to be less than 0.005 (Figure 4E). To measure the errors produced when turning DOR values back to mean expressions, we made inverse prediction models of those three models (See Appendices for details) and tested their performance. As described in Appendices, all inverse prediction models have a fundamental issue in predicting mean expression values for zero DOR, so we excluded those data from performance evaluation. We visualized maximum absolute error (MaxAE) values in addition to MAE values to quantify the prediction performance for data of low DOR. LM exhibited the lowest MAE, scoring less than 0.1 errors in mean expression values on average (Figure 4F), and it performed the best in MaxAE as well (Figure 4G).

Following this, we tested if there is correspondence between DOR and some data attributes unique to individual datasets using a group of datasets that we have named “Mereu2020” as they were obtained in 2020 by Mereu and colleagues [17]. Mereu2020 includes 15 superfamilies where the same sample components were measured across different protocols (e.g., different platforms or different sequencing depths) in order to benchmark scRNA-seq protocols [17], including Chromium V2 (deep), Chromium V2 (shallow), Chromium V2 (sn), Chromium V3, C1HT-medium, C1HT-small, CEL-seq2, Drop-seq, ICELL8, MARS-Seq, Quartz-Seq2, gmcSCRB-seq, ddSEQ, inDrop, and Smart-Seq2; for detailed descriptions, please refer to the original article [17] and URLs of the corresponding webpages on Gene Expression Omnibus that we provide in Methods. Datasets included in Mereu2020 exhibit a wide range of variations in sample sizes and total reads (Figure S6A). When we visualized the coverages of gene expressions (in other words, proportions of non-zero values which are equivalent to 1 − *DOR*), numbers of unique molecular identifier (UMI), and total reads per sample, the datasets with high coverages were found enriched in UMI and read counts (Figure S6B-D). Thus, DOR appears to rejected those metadata attributes, as was discussed in previous studies [12, 13]. Furthermore, we tested whether we could accurately reproduce LMs explaining the intertwinement between DOR and mean expressions when using Mereu2020 datasets (Figure S7A-O). As we showed with the PBMC3k dataset, logistic-transformed mean expression values of all Mereu2020 datasets fitted well to the linear calibration curves, with high coefficients of determination (Figure S8A). We also benchmarked LMs by comparing them with Poisson and NB regression models using those datasets (Figure S7A-O, S8B-E). As detailed in Appendices, the LM demonstrated its capability to function as a calibration curve for DOR and mean expression values across a wide range of datasets. Thus, our analysis indicates that DOR reflects metadata features and per-gene characteristics. Based on the provided examples, we concluded that DOR can be a useful statistic reflecting collective features of scRNA-seq data, including mean expression values and metadata such as sequencing depth information.

### Weighted evaluation function

As stated above, the GRN formation dismisses actual mean expression values of a cell cluster by encoding only the co-occurrence (or co-absence) of gene expressions. To address this issue, we introduced a new metric that can serve as an evaluation function of GRNs instead of *d*^*^. This allows us to assign weights to the similarity of graph structures based on the abundance of gene expressions. To quantify gene expression amounts comparably across different datasets, we used coverage (the presence of non-zero gene expressions, equivalent to 1 − *DOR*), expecting DOR to indirectly reflect the mean expressions of the marker genes forming the edges of the GRNs. With a map *q* : Γ *× X* → ℕ that returns raw gene counts for gene ∀*g* ∈ Γ for sample ∀*x* ∈ *X* (where Γ is the whole set of genes and *X* is the whole set of samples), we formulated the coverage function *Coverage*_[*x*]_ : Γ → ℚ of cell class [*x*] (indicating the coverage value of the given gene *g* in the designated cell class [*x*]) as follows:

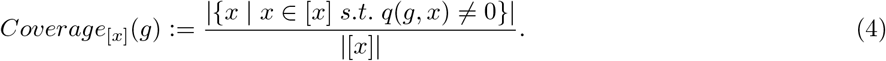

Note that *Coverage*_[*x*]_ relies on *q* only for identifying zeros in raw counts; therefore, any kind of values converted from raw counts by a transformation *ψ* : ℕ→ ℝ such that *ψ*^−1^[{0}] = {0} can be used instead of *q*(*g, x*). For instance, RPM values and log_2_(*RPM* + 1) are accepted (see Appendices for more detailed explanations).

Given that Eq.(1) can also be denoted as Eq.(5), we introduced our new evaluation function, the weighted Hamming quasi-pseudo-metric (WHQPM) *Whqpm*, by multiplying the cardinality of the gene set |*G*| respectively with the coverage values resulting in Eq.(6):

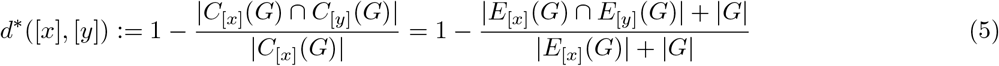

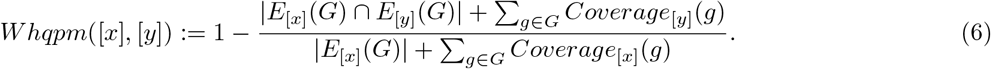

Note that WHQPM cannot be defined if *Coverage*_[*x*]_(*g*) = 0 for all ∀*g* ∈ *G*, and this property of WHQPM prohibits a cell class from being assigned similarity to other cell classes based on totally irrelevant genes exhibiting zero expressions (See also Appendices for detailed explanations).

As WHQPM depends on coverage values, not only biological variations but also technical factors including choices of sequencing pipelines affect the result. If one considers that differences in DOR are also realistic features of the data, WHQPM is available for comparing cell classes across different datasets. Otherwise, optimal transport (OT)-based domain adaptation can mitigate the gap if the experimenter prefers to standardize the various effects that impact on DOR, as described in Figure S9A-H and Appendices.

To demonstrate the benefit of using WHQPM, we first computed GRNs of the clusters 0 through 8 on their top 5 DEGs using the dropout-based binarization technique and the PC algorithm for categorical data (Figure 5A), and then calculated *Whqpm* values to visualize the similarities of the clusters (Figure 5B). Although the PC algorithm for categorical data inferred the exact same GRNs for different clusters in some cases (e.g., the GRNs of the top 5 DEGs of cluster 8, namely *PF4, GNG11, PPBP, SPARC*, and *SDPR*), *Whqpm* distinguished the differences between cluster 8 and the other clusters as it returned non-zero values except for cluster 8 itself. As *d*^*^ returns zero if the subjective cell class has no edges in its GRN, we could resolve this issue with WHQPM.

**Figure 5:**
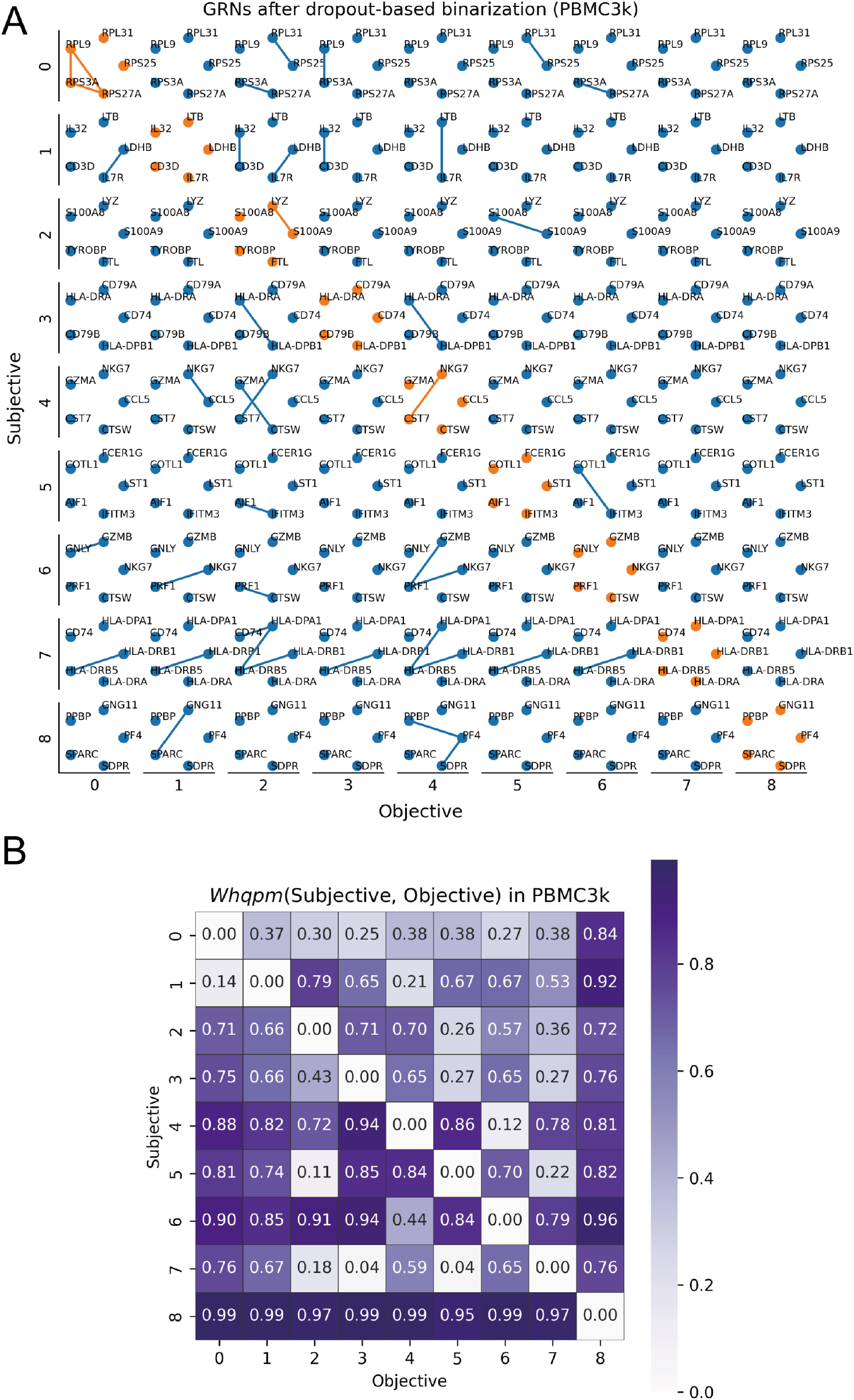
Combination of dropout-based binarization and WHQPM. **A**: GRNs of the clusters in PBMC3k generated with dropout-based binarization and the PC algorithm for categorical data. GRNs in a row share the same set of genes (DEGs of the subjective clusters) selected for the vertex sets. **B**: The *Whqpm* values based on the GRNs of the top 5 DEGs generated after dropout-based binarization.

## Discussion

In general, scRNA-seq data processing is driven by statistical, geometrical, and information-theoretical approaches, even though the results from these algorithms are validated by their ability to recite the storyline of biology. In other words, the specific details of algorithms are not necessarily interesting as long as the results make biological sense. Therefore, heuristics is often valued over theoretical rigor. As scRNA-seq data accumulate at an accelerating rate, and despite their sensitivity to fluctuations in surrounding conditions, we believe that a framework that can handle scRNA-seq data in a tentative but comparable format will help balance context-dependency and generalization, uncovering universal truth yet to be unveiled.

We previously designed the GRN-based definition of cellular identities and the metric *d*^*^ to quantify their similarity levels. However, our method included several impracticalities, as described earlier in this article. Therefore, we proposed a series of solutions aimed at improving functionality. Additionally, we launched a Python package, GRNet (pronounced garnet), to provide a platform for our proposed concepts. While promising results have been observed in a limited number of datasets, broader validation and discussion is still required.

## Methods

### GRNet Implementations

#### GO term-assisted gene selection using Jaccard Index

The Jaccard Index of two sets *A, B* is defined as:

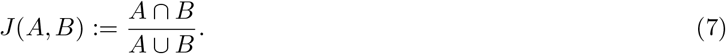

We expanded this definition to pairwise comparisons of multiple elements by forming a matrix where each element is the corresponding Jaccard Index. We named this matrix the Jaccard index matrix (JIM). As an example, the element in the *i*-th row and *j*-th column (where *i, j, k* ∈ ℕ and *i* ≤ *k, j* ≤ *k*) is defined as follows when a JIM of sets *X*_1_, *X*_2_, · · ·, *X*_*k*_ is introduced:

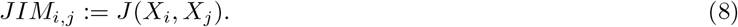

When a collection of genes *g*_1_, · · ·, *g*_*k*_ collectively explain a certain type of cells, and they are tagged with respective sets of GO terms *G*_1_, · · ·, *G*_*k*_, we considered *min*_*i,j*∈{1···*k*}_*J* (*G*_*i*_, *G*_*j*_) as a threshold of biological correspondence to the type of cells. For example, let *G*_*k*+1_, *G*_*k*+2_ be the sets of GO terms tagged with *g*_*k*+1_ and *g*_*k*+2_ (*g*_*k*+1_, *g*_*k*+2_ ∉ {*g*_1_, · · ·, *g*_*k*_}); then a new gene *g*_*k*+1_ would be important for the type of cells if *min*_*i*∈{1···*k*}_*J* (*G*_*i*_, *G*_*k*+1_) is less than *min*_*i,j*∈{1···*k*}_*J* (*G*_*i*_, *G*_*j*_), and *g*_*k*+2_ would be irresponsible if *min*_*i*∈{1···*k*}_*J* (*G*_*i*_, *G*_*k*+2_) is greater than *min*_*i,j*∈{1···*k*}_*J* (*G*_*i*_, *G*_*j*_). Under those rules, we implemented a search for important markers from genes tagged with GO terms in ⋂_*i*∈{1,···, *k*}_ *G*_*i*_.

For detailed implementation, we calculated the JIM of the related GO terms of given seed markers. We used mygene.py [18] to query the GO database, and Numpy [19] to calculate JIM.

#### GRNs and the evaluation function

Following our previous report [3], we implemented a correlation-baed PC algorithm for GRN formation and the evaluation function *d*^*^ for similarity of GRN structures using Numpy, Pandas [20], and pgmpy. Additionally, we implemented dropout-based binarization, a chi-squared test-based PC algorithm, and WHQPM accordingly.

### scRNA-seq data analysis

#### Dataset List

The scRNA-seq data we used in this research were publicly available from the following online resources:

- PBMC3k: https://support.10xgenomics.com/single-cell-gene-expression/datasets/1.1.0/pbmc3k
- Mereu2020: https://www.ncbi.nlm.nih.gov/geo/query/acc.cgi?acc=GSE133549
  – Chromium V2 (deep): https://www.ncbi.nlm.nih.gov/geo/query/acc.cgi?acc=GSE133535
  – Chromium V2 (shallow): https://www.ncbi.nlm.nih.gov/geo/query/acc.cgi?acc=GSE133536
  – Chromium V2 (sn): https://www.ncbi.nlm.nih.gov/geo/query/acc.cgi?acc=GSE133546
  – Chromium V3: https://www.ncbi.nlm.nih.gov/geo/query/acc.cgi?acc=GSE141469
  – C1HT-medium: https://www.ncbi.nlm.nih.gov/geo/query/acc.cgi?acc=GSE133537
  – C1HT-small: https://www.ncbi.nlm.nih.gov/geo/query/acc.cgi?acc=GSE133538
  – CEL-seq2: https://www.ncbi.nlm.nih.gov/geo/query/acc.cgi?acc=GSE133539
  – Drop-seq: https://www.ncbi.nlm.nih.gov/geo/query/acc.cgi?acc=GSE133540
  – ICELL8: https://www.ncbi.nlm.nih.gov/geo/query/acc.cgi?acc=GSE133541
  – MARS-Seq: https://www.ncbi.nlm.nih.gov/geo/query/acc.cgi?acc=GSE133542
  – Quartz-Seq2: https://www.ncbi.nlm.nih.gov/geo/query/acc.cgi?acc=GSE133543
  – gmcSCRB-seq: https://www.ncbi.nlm.nih.gov/geo/query/acc.cgi?acc=GSE133544
  – ddSEQ: https://www.ncbi.nlm.nih.gov/geo/query/acc.cgi?acc=GSE133547
  – inDrop: https://www.ncbi.nlm.nih.gov/geo/query/acc.cgi?acc=GSE133548
  – Smart-Seq2: https://www.ncbi.nlm.nih.gov/geo/query/acc.cgi?acc=GSE133545

#### Preprocessing, dimensionality reduction, and visualization

We performed data preprocessing, dimensionality reduction, and data visualization of the scRNA-seq datasets using Python packages (including Scanpy [21], Polars [22], Pandas, Numpy, Matplotlib [23], and Seaborn [24]).

#### Clustering and DEA

We performed leiden clustering, and DEA using Scanpy.

#### Multiclass classification GBDT model

We randomly split the PBMC3k data into training, validation, and test data (3:1:1). Using the training and validation data, we created a GBDT model minimizing the multiclass-logarithmic loss function using LightGBM’s framework [25]. We implemented the model with a wrapper in Optuna [26] to automatically tune the hyperparameters. The models performance was tested with ROC curves and PR curves using Scikit-learn [27] and Matplotlib. We also visualized the feature importance values implemented in LightGBM. SHAP scores were calculated and visualized with the Shap Python package [11, 28].

#### GO analysis

We performed the GO analysis using gprofiler2 [29], and visualized the results with Matplotlib and Seaborn.

#### Statistical models of DOR and the benchmarking

For LM, we optimized *b* of the calibration curve by minimizing the MSE between *DOR* and 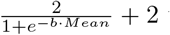 with AdaGrad. We implemented LM and plotting functions with AnnData, Matplotlib, Numpy, Pandas, and PyTorch [30]. We implemented Poisson regression models with Statsmodels [31]. For NB regression models, we built them on Statsmodels and optimized hyperparameters using Optuna.

#### OT-based coverage standardization

We made OT-based domain adaptation models using the EMDTransport class of POT [32] with the squared Euclidean cost. The results were visualized with Matplotlib, Numpy, Pandas, and Seaborn.

### Other visualizations

#### Alluvial plot and Venn diagram for GO terms

The glial markers were selected based on review articles, and the tagged GO terms were queried using mygene.py. Then, all gene symbols subscribed with each GO terms were queried again. The alluvial plot was created with Matplotlib, Numpy, and Pandas, and the Venn diagram was visualized with Matplotlib-Venn [33].

## Code availability

GRNet and the analysis codes are available on GitHub at https://github.com/yo-aka-gene/grnet. Online documentation for GRNet is also provided at https://grnet.readthedocs.io.

## Author contributions

**Conceptualization** YO

**Methodology** YO

**Implementation** YO

**Investigation** YO

**Visualization** YO

**Funding acquisition** YO, YK, HO

**Project administration** YO, YK, HO

**Supervision** HO

**Senior author** YK

**Original draft** YO

**Editing** YK, HO

## Acknowledgements

This work was supported by the Keio University Medical Science Fund (to YO). We are grateful to Dr. Hans Dijkstra (Fujita Health University) for critical reading of the manuscript.

## Abbreviations

AP: average precision
AUC: area under the curve
DEA: differential expression analysis
DEG: differentially expressed gene
DOR: dropout rate
GBDT: gradient boosting decision tree
GO: gene ontology
GRN: gene regulatory network
HVG: highly variable gene
JIM: Jaccard index matrix
LM: logistic model
MAE: mean absolute error
MaxAE: maximum absolute error
ML: machine learning
MMHC: max-min hill-climbing
MSE: mean squared error
NB: negative binomial
OT: optimal transport
OvR: one-versus-rest
PC: Peter and Clark
PCA: principal component analysis
PR: precision-recall
QC: quality control
ROC: receiver operating characteristic
RPM: reads per million
scRNA-seq: single-cell RNA-sequencing
SHAP: Shapley additive explanations
TSVD: truncated singular value decomposition
UMAP: uniform manifold approximation and projection
UMI: unique molecular identifier
WHQPM: weighted Hamming quasi-pseudo-metric

## Appendices

### Supplemental Information

#### Cell clusters vs. cells classes

A cell cluster refers to a stochastic chunk of samples neighbouring each other in the data space or specific feature space designed via the data analysis. Hence, cell clusters can be defined without introducing GRNs or eigen-cascades. In contrast, a cell class refers to a set of samples that can be treated identically, with their biological features characterized by eigen-cascades. In this context, what is identical can be defined by a clustering function (i.e., a mapping between samples and the clusters to which they belong) of choice. Therefore, the term cell class emphasizes the randomness of a cell cluster and is always tagged with the concept of eigen-cascades. On the other hand, we use cell cluster to refer some data-driven groups of cells in a more neutral nuance.

#### The vertex set *V*_[*x*]_(*G*), the edge set *E*_[*x*]_(*G*), and the eigen-cascades *C*_[*x*]_(*G*) of a cell class [*x*]

A graph (*V, E*) is a pair of a vertex set *V* and an edge set *E*. To create a graph of a set of genes *G* based on the statistical dependencies among the genes, the vertex set *V* (*G*) and the edgeset *E*(*G*) are introduced as follows:

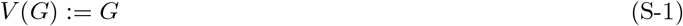

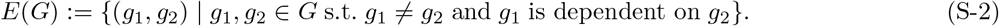

It is important to note that (*V* (*G*), *E*(*G*)) itself refers to a multigraph and we need to equate (*g*_1_, *g*_2_) and (*g*_2_, *g*_1_) to create GRNs (undirected simple graphs). In the previous article, we introduced asymmetrical notations for edge sets to build our theory from causal networks and relaxed the restriction by replacing the causalities with the statistical dependencies. In this section, we adopted the same style for consistency, even though the statistical dependency of two variables is symmetrical.

We also introduced a notation 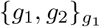 where ∀*g*_1_, *g*_2_ ∈ *G* such that *g*_1_ is dependent on *g*_2_, representing an indexed set with the index *g*_1_ ∈ {*g*_1_, *g*_2_} (note that the symmetry of statistical dependency induces symmetry 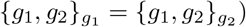. We named such indexed sets cascades. The inclusion of *g*_1_ being dependent on itself leads to the expression 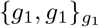 ; however, mathematical set rules simplify this to 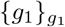. Notably, these singleton sets are also considered valid representations of cascades. The eigen-cascades of a cell class [*x*] are the cascades that can be inferred from [*x*], which is the cell class that a sample *x* belongs. As all cascades are either singletons or sets of two elements, the set of eigen-cascades *C*_[*x*]_(*G*) of [*x*] equals to the union of *V*_[*x*]_(*G*) and *E*_[*x*]_(*G*), defined as follows:

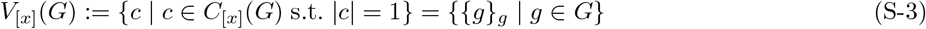

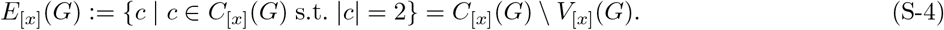

Juxtaposing the given definitions from Eq. (S-1) to Eq. (S-4), *V*_[*x*]_(*G*) and *V*_*x*_(*G*) as well as *E*_[*x*]_(*G*) and *E*_*x*_(*G*) have similar formulations. More precisely, *V*_[*x*]_(*G*) is isomorphic to *G* and *E*_[*x*]_(*G*) can be isomorphic to some edge set (e.g., a map 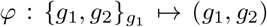 is an isomorphism and *φ*(*E*_[*x*]_(*G*)) is a valid edge set). Accordingly, a graph (*G, φ*(*E*_[*x*]_(*G*))) can be formed referring to the eigen-cascades, and the pair of *C*_[*x*]_(*G*) and (*G, φ*(*E*_[*x*]_(*G*))) show one-to-one correspondence. Concisely, *V*_[*x*]_(*G*) and *E*_[*x*]_(*G*) can be regarded as the vertex set and the edge set.

It is important to note that the mathematical discussions to extend the notion of cascades to eigen-cascades of cell classes include various notations, operations, and hypotheses that can sound confusing or non-trivial. In this article, we simply introduced the notations of [*x*] and *C*_[*x*]_(*G*) aiming to shortcut the redundant part. We recommend referring to the original article for readers interested in the detailed explanations.

#### The logistic model of DOR

Although the mean expression values and the DOR values show a non-linear relationship, we tried to find a simple non-linear conversion that enables the mean expression values to linearly fit to the DOR values. We leveraged the logistic transformation (i.e., inverse logit transformation) so that the transformed values become positive and less than 1. Introducing *u* to be the variable for the transformed values as follows, we tried to fit the parameter *b* (*b >* 0) so that *DOR* ∼ −2*u* + 2 by minimizing the MSE:

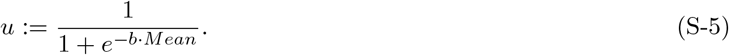

Note that the minimal value of *u* is 0.5 (when *Mean* = 0) and the following equation holds for the limit:

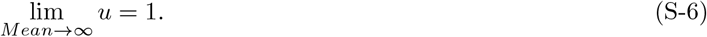

In addition to those properties, the range of DOR is 0 ≤ *DOR* ≤ 1, and they are the reasons why we designed the fitting line as *DOR* ∼ −2*u* + 2. Consequently, the final form of the fitting curve can be denoted as follows:

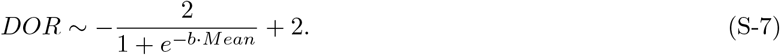

#### Inverse predictions of LM, Poisson regression, and NB regression

The inverse prediction of LM can be formulated as follows by solving Eq. (S-7) iff *DOR* ≠ 0:

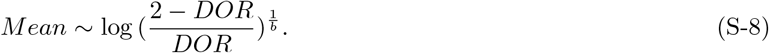

As we used the logarithmic link function for both the Poisson regression models and the NB regression models, those models can be formulated as follows:

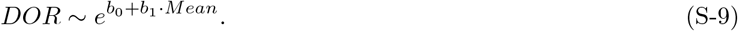

Therefore, the inverse predictions of those models can be formulated as follows by solving Eq. (S-9) iff *DOR* ≠ 0:

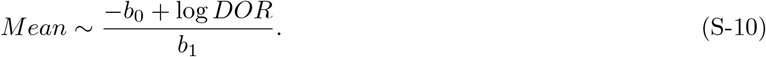

It is noteworthy that the *Mean* value diverges into infinity as *DOR* approaches zero, regardless of the model choice. Therefore, the performance comparison of the inverse prediction models was conducted while excluding data with zero *DOR* values to prevent metrics such as MAE from diverging into infinity. This ensures that the results do not indicate that all models perform equally poorly by generating infinite prediction errors on average.

#### Benchmarking DOR regression models with Mereu2020 datasets

For benchmarking DOR regression models (namely LM, Poisson regression model, and NB regression model), we used Mereu2020 dataset families so that we could evaluate the performance of those models when the data were acquired in various sequencing protocols. As we did with PBMC3k dataset, we created LMs, Poisson regression models, and NB regression models for respective datasets (Figure S7A-O). Then we measured MSE (Figure S8B), MAE (Figure S8C), MAE of the inverse-prediction models (Figure S8D), and MaxAE of the inverse-prediction models (Figure S8E) to evaluate their performance.

Unlike the result from the PBMC3k dataset, LM underperformed Poission and NB regression models in most cases (except for Chromium V2 (sn), Chromium V3, C1HT-medium, and C1HT-small) in terms of MSE (Figure S8B). Similar results were reproduced in terms of MAE even though all models scored quite small MAE values (Figure S8C). Although LM slightly underperformed the two conventional regression models in explaining the relationship between mean expressions and DOR values in general cases, LM showed stable performance in the inverse-prediction tasks scoring less than 0.1 MAE in all cases (Figure S8D), and even outperformed Poisson and NB regression models in terms of MaxAE (Figure S8E), suggesting that LM performed better than the other two models at low DOR values. These results indicated that LM models work as calibration curves that can transform DOR and mean expression values interchangeably in wide ranges of those input values, while the conventional regression models have more weight on domains where samples densely distribute (regularly high-DOR and low-mean-expression areas) to outperform LM in terms of MSE, MAE, and MAE of the inverse prediction in general.

#### The preimage and general scRNA-seq data transformation

Let ∀*A, B, C* be sets such that *C* ⊂ *B*, and let *f* : *A* → *B* be a map. The preimage of *C* under *f* (denoted as *f* ^−1^[*C*]) is defined as follows:

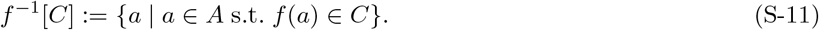

Therefore, a function *ψ* : ℕ → ℝ such that *ψ*^−1^[{0}] = {0} returns zero if and only if the input value is zero. This property is essential for *Coverage*_[*x*]_ to identify non-zero values.

Let *ψ*_*R*_ : ℕ → ℚ as the RPM transformation from the raw counts, and let *ψ*_*L*_ : ℚ → ℝ be the transformation from RPM values to log_*k*_ (*RPM* + 1) for all *k >* 0. It is trivial that 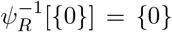 and 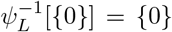 hold. Accordingly, the conversion from raw counts to log_*k*_ (*RPM* + 1) values are *ψ*_*L*_ *° ψ*_*R*_, and the following equation holds:

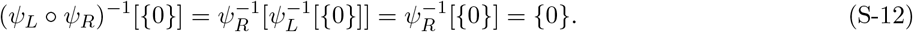

#### An example where WHQPM is undefined

Looking at the definition of *Whqpm*, it is clear that the formula holds iff |*E*_[*x*]_(*G*)| + Σ_*g*∈*G*_ *Coverage*_[*x*]_(*g*) ≠ 0. Given that both |*E*_[*x*]_(*G*)| and Σ_*g*∈*G*_ *Coverage*_[*x*]_(*g*) are non-negative, *Whqpm* becomes undefined when these two terms are both zero. Such a condition is readily satisfied when the GRN of the subjective cell class [*x*] is characterized by genes completely absent in [*x*], exhibiting zero coverages (i.e., DOR values of those genes are all 1). In such a case, not only is Σ_*g*∈*G*_ *Coverage*_[*x*]_(*g*) = 0, but also |*E*_[*x*]_(*G*)| becomes 0 (otherwise ∀*e* ∈ |*E*_[*x*]_(*G*) is undefined). The status of |*E*_[*x*]_(*G*)| varies depending on the choice of the GRN formation algorithm, in other words, computational methods to assign edges among vertexes. While the correlation-based method and the chi-squared test-based method coupled with the dropout-based binarization are not applicable for the case where the mean expression values are all zero and all samples in [*x*] are negative for all genes, Fisher’s exact test is an available option for the algorithm to compute the edges. However, the null hypotheses fail to be rejected in those setting resulting in |*E*_[*x*]_(*G*)| = 0.

Consequently, *Whqpm* is undefined when the GRN of [*x*] is formed with genes such that Σ_*g*∈*G*_ *Coverage*_[*x*]_(*g*) = 0. By design, WHQPM prevents ill-defined characterization of the similarity between two cell classes by staying undefined when the subjective cell class is represented exclusively with uncharacteristic genes.

#### OT-based domain adaptation for coverage standardization

To demonstrate the OT-based coverage standardization across different sequencing platforms, we consider standardizing the distributions of coverage values using C1HT-medium dataset and the inDrop dataset from Mereu2020. Let the distribution of inDrop be the source domain, which is to be standardized, and that of C1HT-medium be the target domain, referred to as the goal of the standardization procedure. As visualized by the inferred distributions upon kernel density estimation (Figure S6B and S9A), inDrop is abundant in genes with zero counts for all samples, and here we aim to correct it using C1HT-medium as a reference.

After randomly undersampling 10,000 genes commonly measured in the both datasets (to reduce computational costs), we performed OT on empirical distributions using earth mover’s distance with squared Euclidean cost (Figure S9B). As the line indicating the coupling of the source data and the transformed data were orderly aligned, the OT model mapped the coverage values in the inDrop dataset to the data distribution of C1HT-medium while preserving the relative order of the original values.

This technique can be extended to adjust for differences in coverage values among clusters in the same dataset. As the definition of coverage depends on the sample sizes of the cell classes, the distributions of coverage values vary depending on the cluster sizes; large clusters have relatively smooth and almost continuous distributions, while small clusters have discrete and stepwise distributions (Figure S9C-D). Furthermore, clusters might exhibit drastic differences in coverage distributions due to geometrical reasons originating from the stochasticity of dropout events. When we tried to standardize the distribution of cluster 5 in PBMC3k to the distribution of cluster 8 (Figure S9E), the same OT model successfully transformed the source distribution into the target one (Figure S9F). To quantify how much OT models decayed the biological characteristics of the source cluster by transformation, we calculated the correlation coefficients of coverage values per gene among the source, transformed, and target distributions (Figure S9G). As the transformed cluster 5 showed a high correlation with the original cluster 5 while the coefficient with cluster 8 was only 0.45, indicating poor correlation, it suggeested that the OT model did not alter the biological features of cluster 5 to resemble those of cluster 8. On the other hand, the same of heatmap regarding the OT model transforming coverage values of inDrop into the distribution of C1HT-medium indicated the model could make samples of the same biological profiles more similar to each other (Figure S9H). Therefore, we conclude that the OT-based domain adaptation model could tone down the differences in coverage values by suppressing the gaps derived from artifacts while preserving the biological uniqueness of the samples.

Although our model performed well in moving the finer distribution (cluster 5 in PBMC3k) to the coarser one (cluster 8), we would not recommend performing the opposite because the coupling practically fails to form a well-defined map that returns the same values for the same inputs. For example, cluster 8 is formed by eleven cells, and the genes can exhibit twelve patterns of coverage values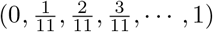. Since the number of genes is shared between clusters 8 and 5, and the OT model finds the best one-to-one correspondence minimizing the cost, there exists a pair of genes that show the exact same coverage values in cluster 8 but get transformed into different values because a cluster of a larger size can show more variation in coverage values than twelve. To avoid ill-defined operations, we recommend applying our method to adjust coverage values by referring to the smaller cell class.

## Supplemental Figures

**Figure S1:**
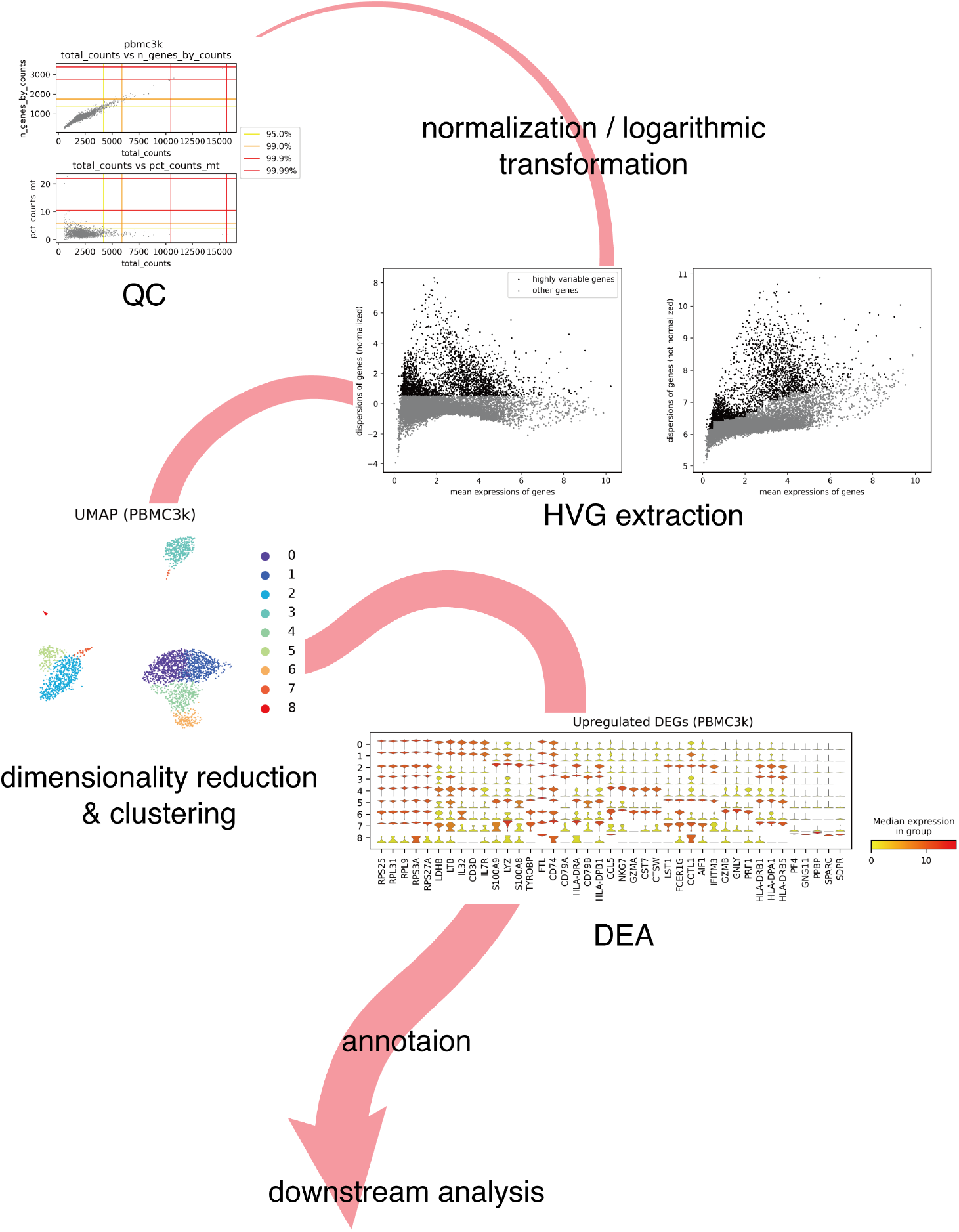
Standard workflow of scRNA-seq analysis. A scheme that shows a standard process of scRNA-seq data analysis. The plots were generated during the analysis of PBMC3k.

**Figure S2:**
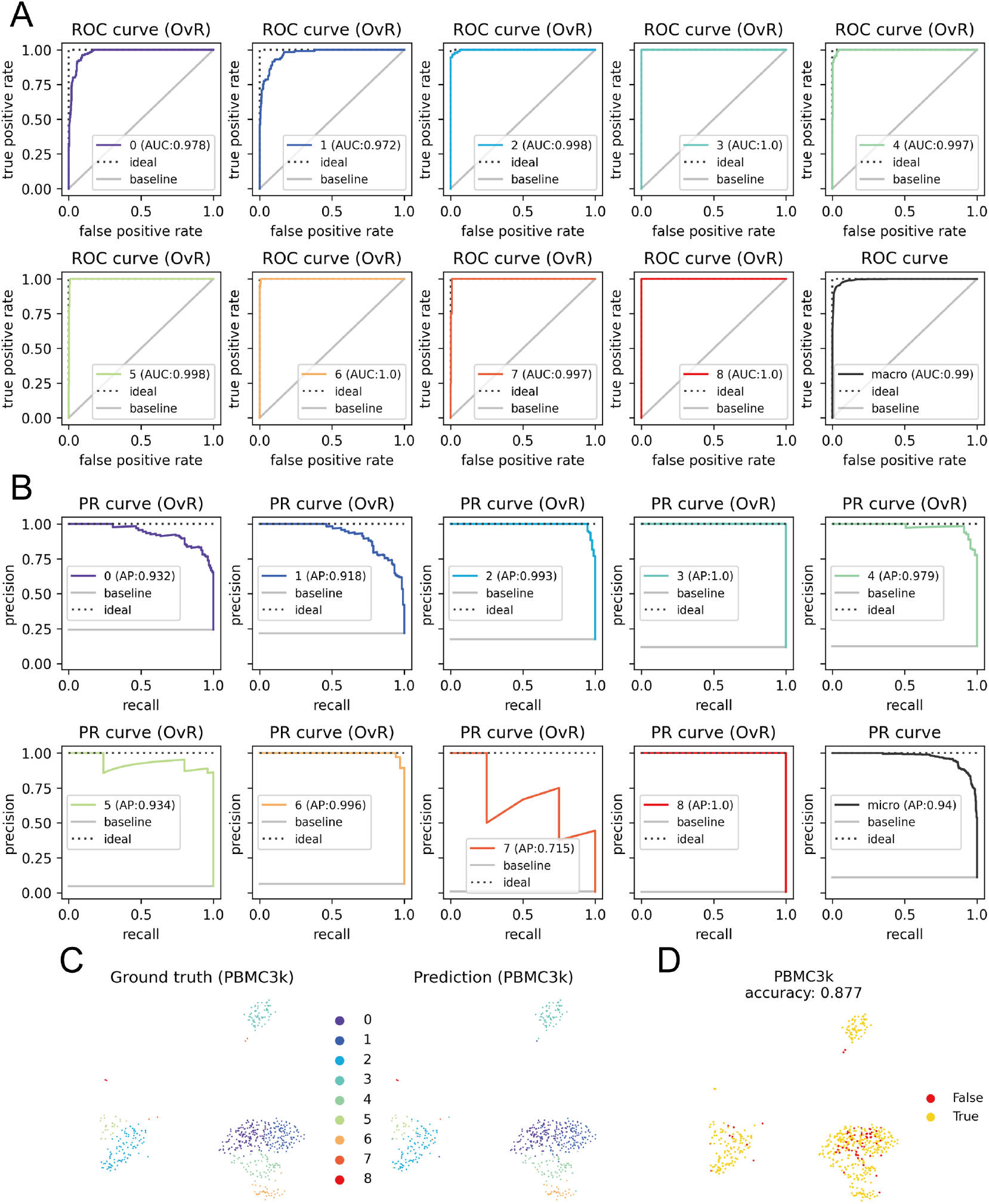
The performance of the GBDT model. **A**: The OvR ROC curves of the clusters 0∼8 and their macro average. **B**: The OvR PR curves of the clusters 0∼8 and their micro average. **C**: A pairwise comparison of the ground-true labels and the predictions. **D**: A UMAP visualization of the prediction accuracy.

**Figure S3:**
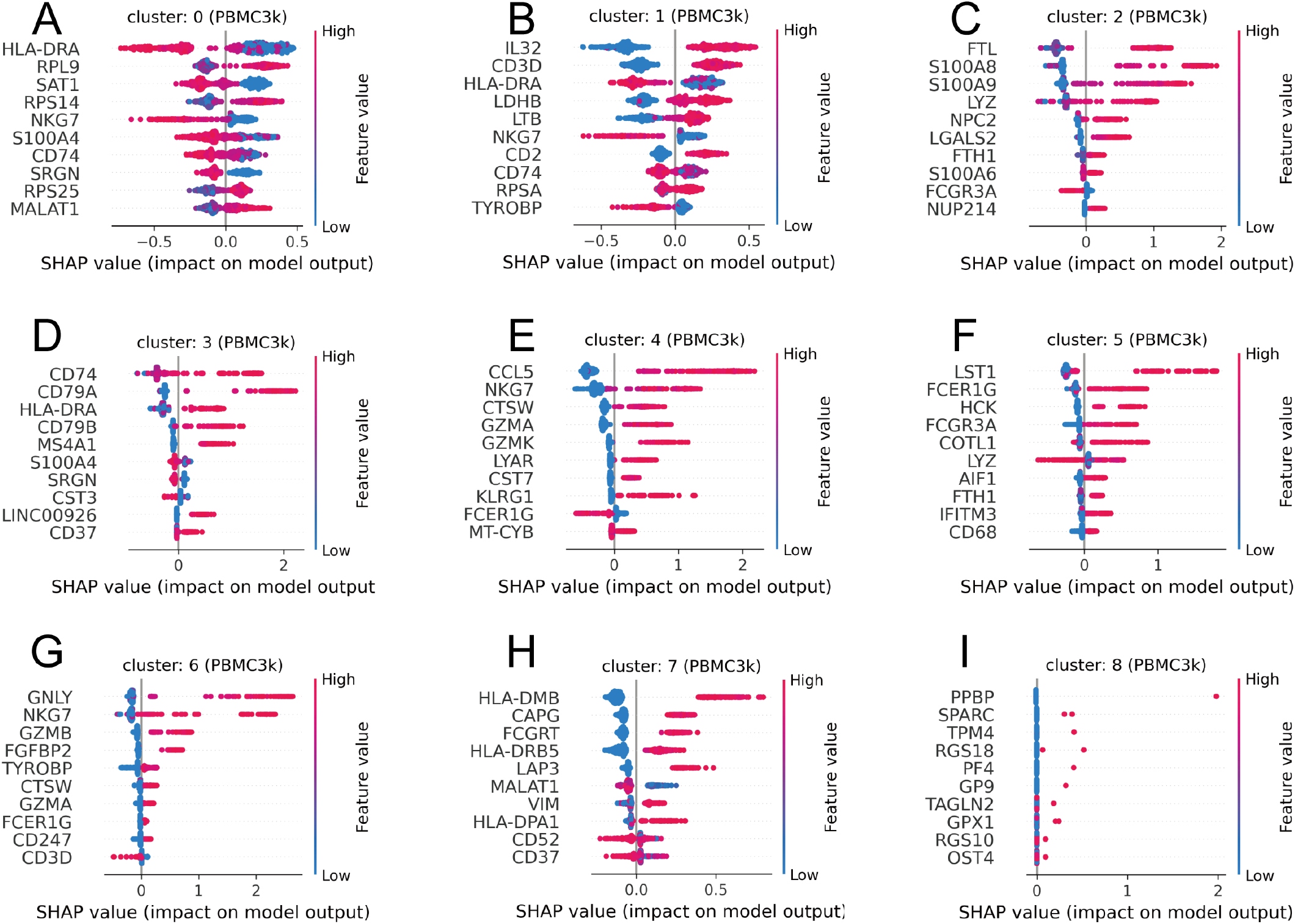
The top 10 features of the clusters in PBMC3k. **A-I**: The top 10 features (based on SHAP values) for cluster 0∼8 respectively.

**Figure S4:**
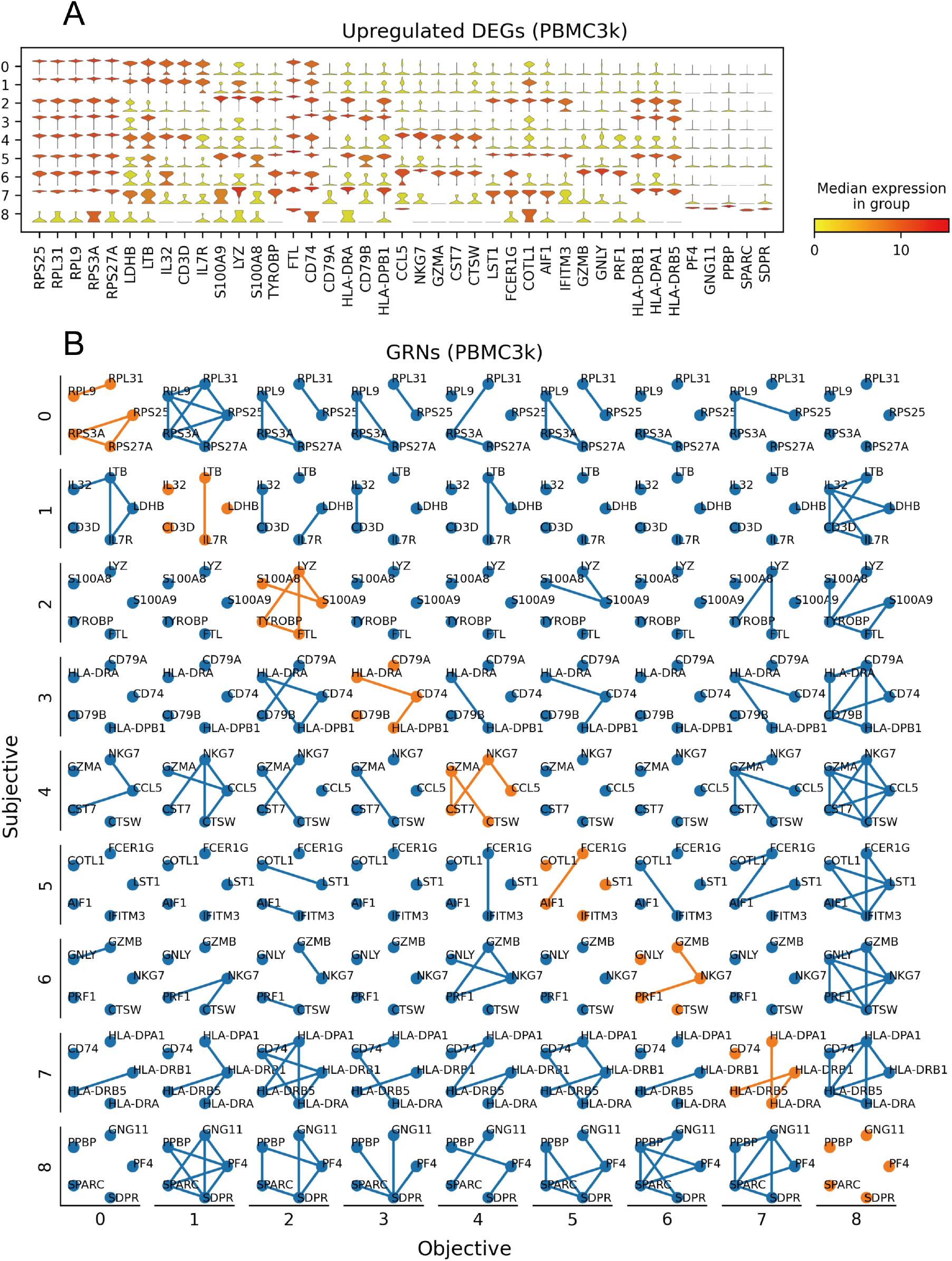
The top 5 DEGs and the GRNs of the clusters in PBMC3k. **A**: Violin plot of DEGs of the clusters (top 5 for each). **B**: GRNs of the clusters. GRNs in a row share the same set of genes (DEGs of the subjective clusters) selected for the vertex sets.

**Figure S5:**
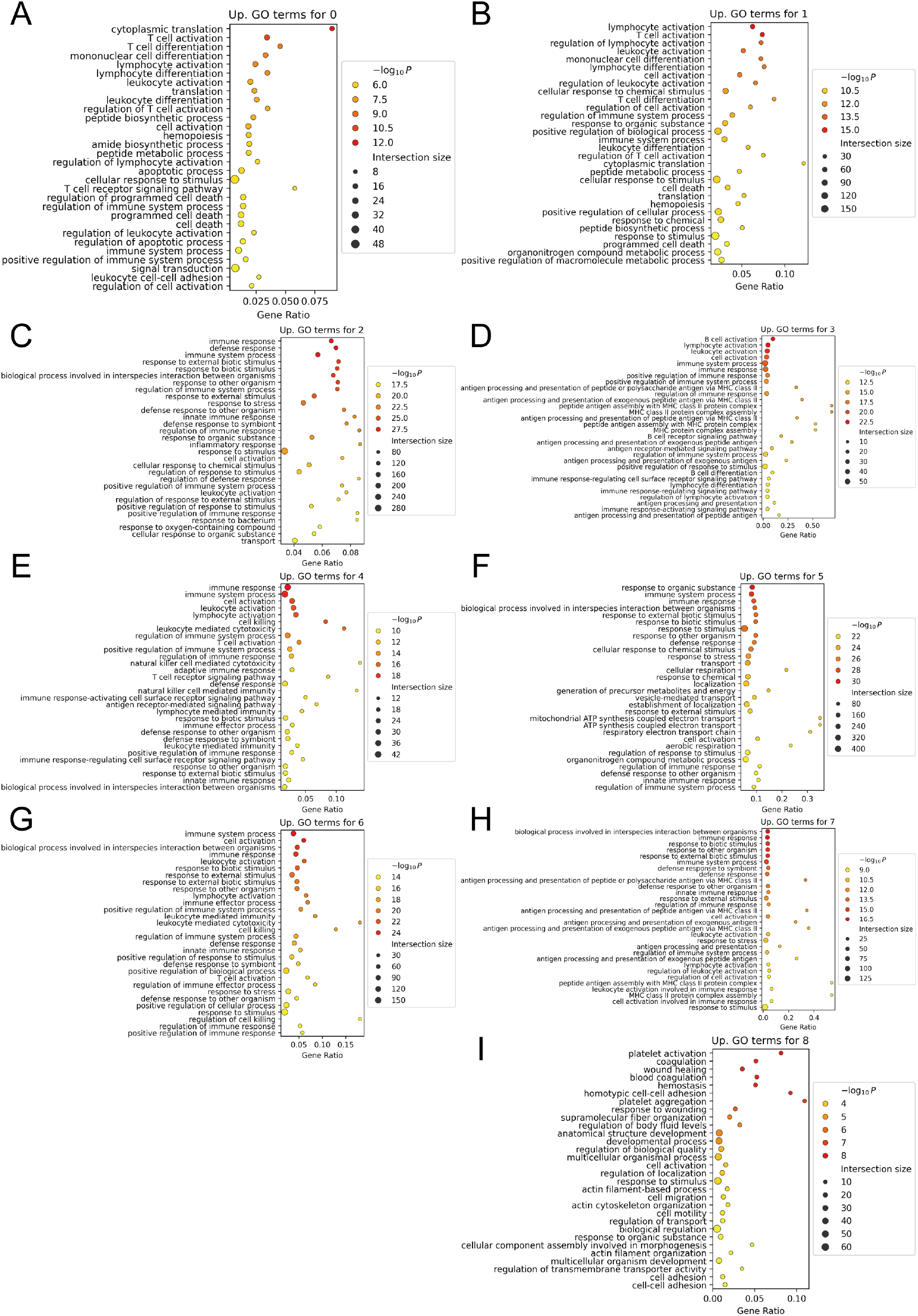
The top 30 upregulated GO terms of the clusters in PBMC3k. **A-I**: The top 30 GO terms for cluster 0∼8 respectively.

**Figure S6:**
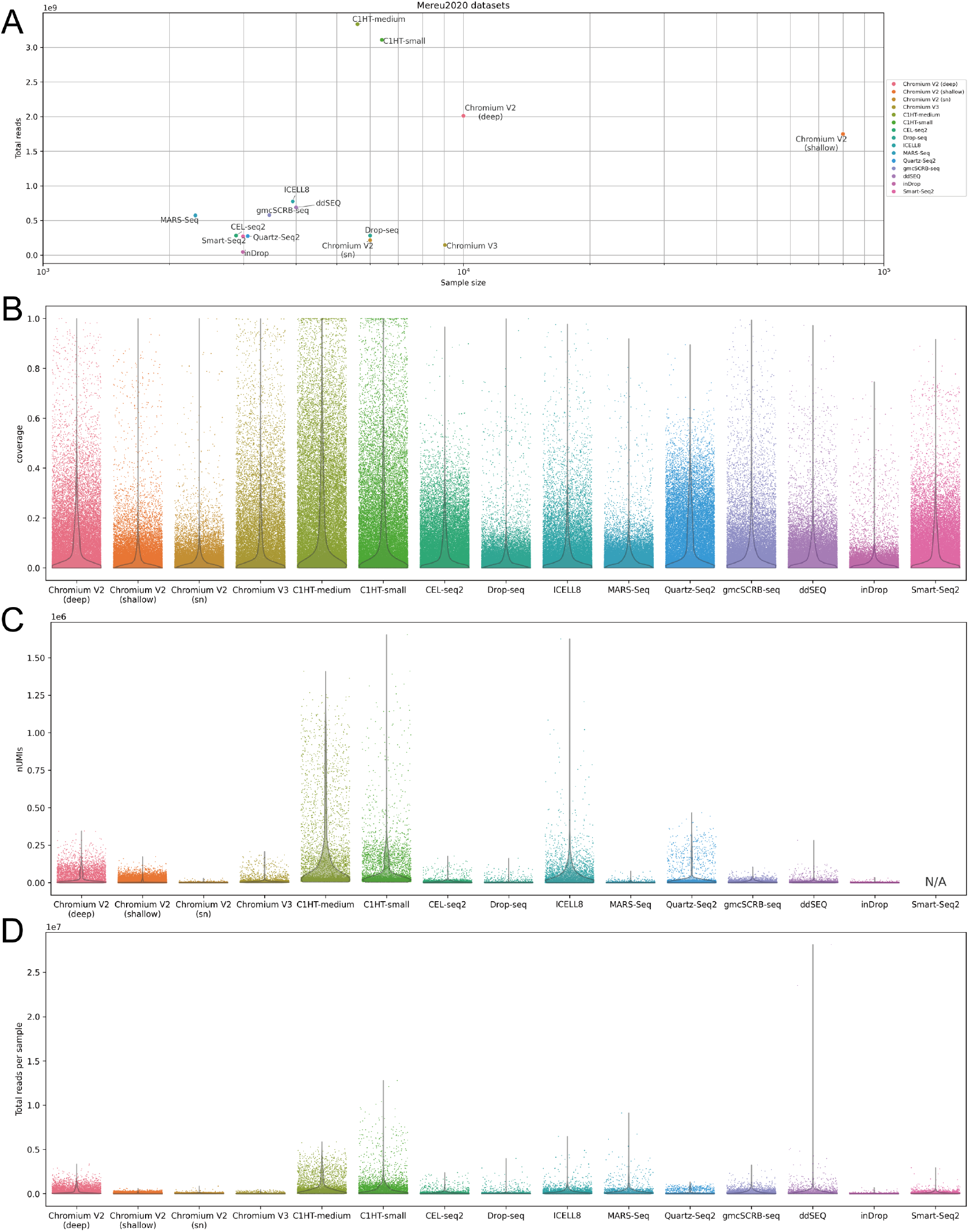
Mereu2020 datasets. **A**: A scatter plot regarding the sample sizes and the total reads of the superfamilies in Mereu2020. **B**: Coverages in the superfamilies in Mereu2020. The dots of the strip plots refer to the coverage values of genes in each dataset. **C**: Numbers of UMIs (nUMIs) in Mereu2020. As SmartSeq2 is not UMI-based, data were not available. **D**: Total reads per sample in Mereu2020 datasets.

**Figure S7:**
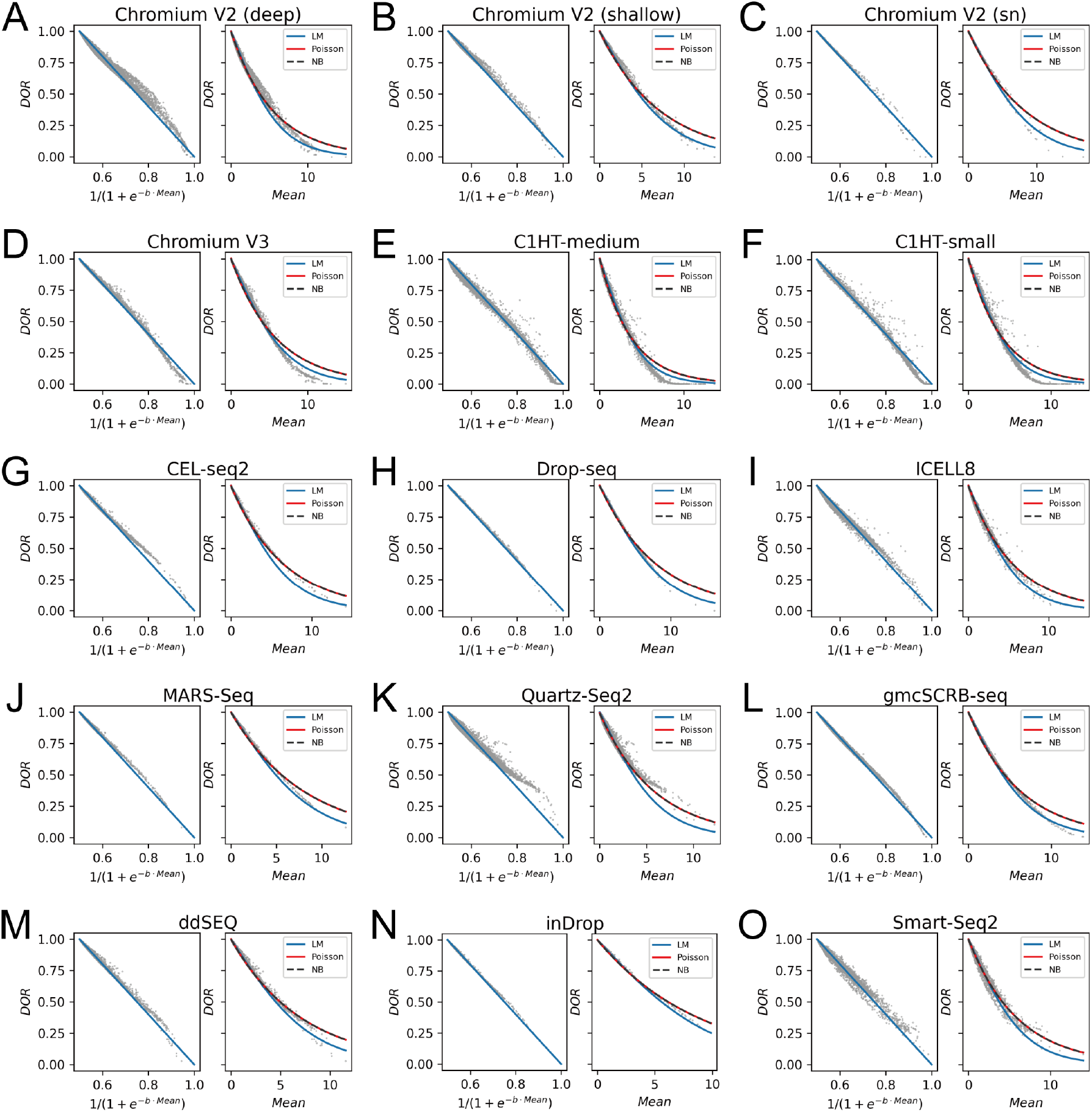
Regression models of DOR in Mereu2020 datasets. **A-O**: A scatter plot regarding the sample sizes and the total reads of the superfamilies in Mereu2020.

**Figure S8:**
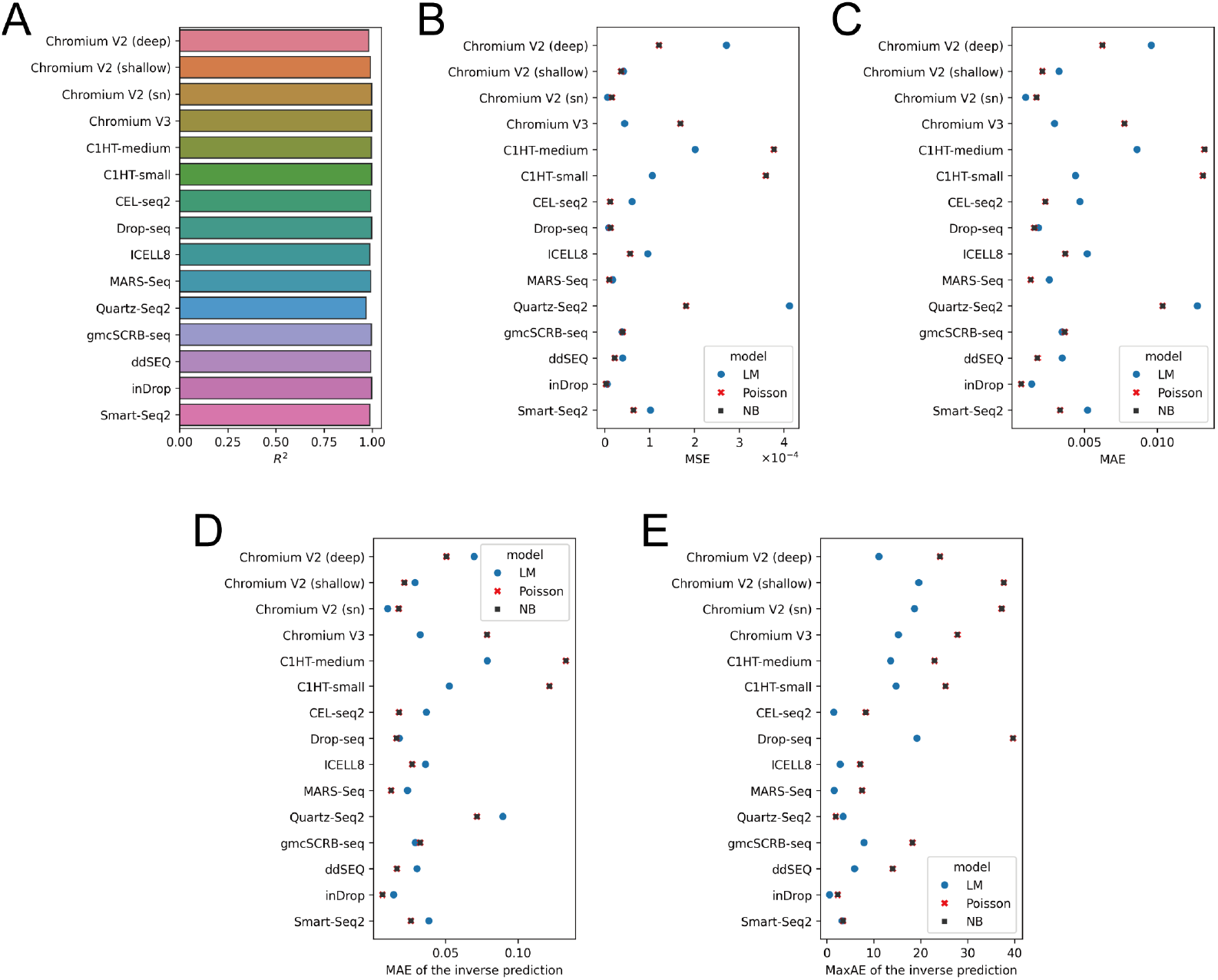
Performance metrics for regression models of DOR in Mereu2020 datasets. **A**: The coefficients of determination (*R*^2^) of logistic-transformed mean expression values to the linear calibration curve of DOR. **B**: Performance comparison of LM, Poisson, and NB with MSE values. **C**: Performance comparison of LM, Poisson, and NB with MAE values. **D**: Performance comparison of the inverse predictions of LM, Poisson, and NB with MAE values. **E**: Performance comparison of the inverse predictions of LM, Poisson, and NB with MaxAE values.

**Figure S9:**
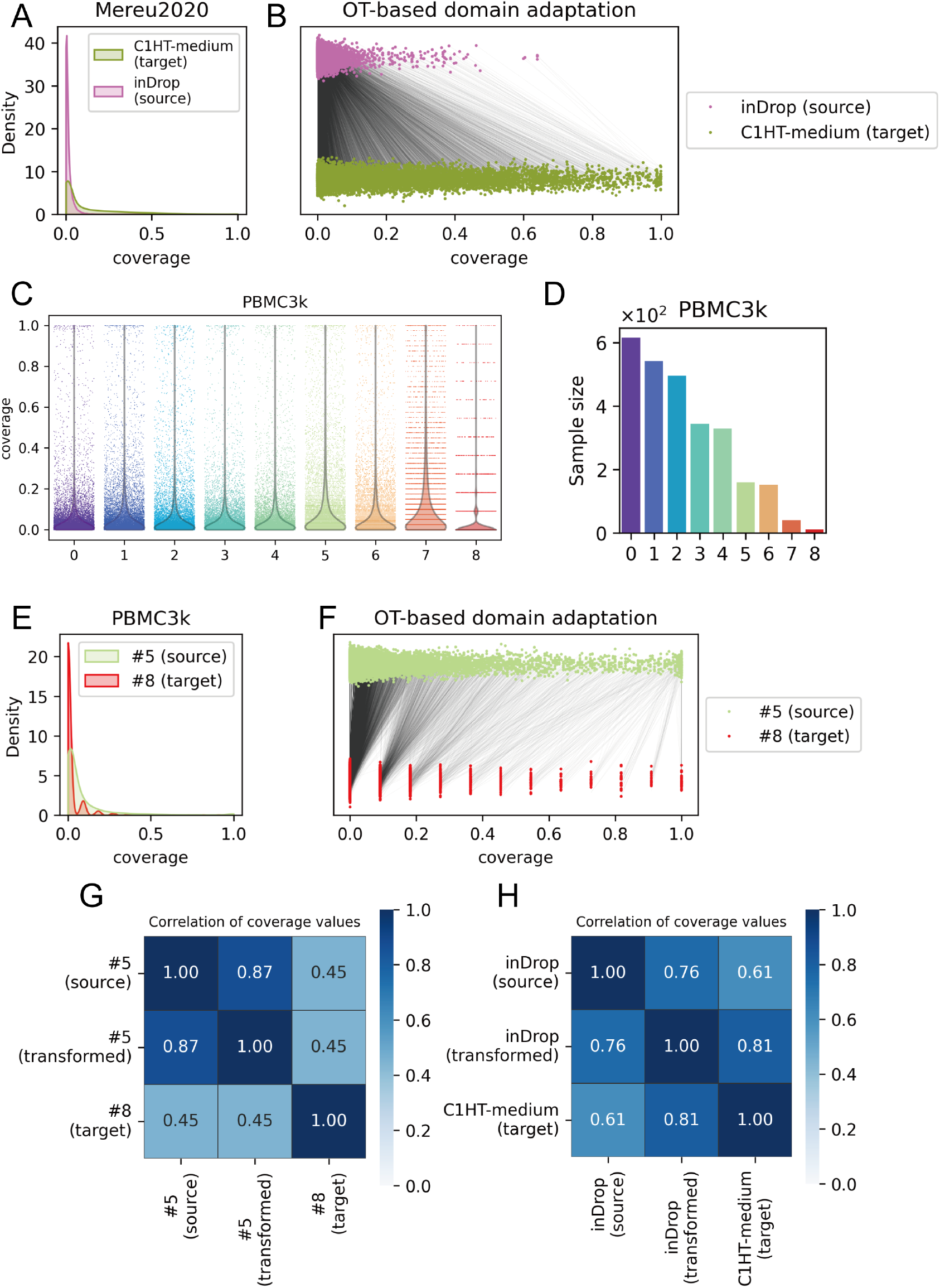
OT-based domain adaptation on empirical distributions of coverage values. **A**: Kernel density estimation of the coverage values in C1HT-medium and inDrop. **B**: Results of OT-based domain adaptation converting the source distribution (inDrop) to the target distribution (C1HT-medium). The lines indicate the pairing computed with OT. **C**: Coverage per cluster in PBMC3k. **D**: Sample size per cluster in PBMC3k. **E**: Kernel density estimation of the coverage values in clusters 5 and 8 of PBMC3k. **F**: Results of OT-based domain adaptation converting the source distribution (cluster 5) to the target distribution (cluster 8). **G**: Correlation coefficients of the coverage values in the source distribution (cluster 5), the transformed distribution (cluster 5 mapped by the OT model), and the target distribution (cluster 8). **H**: Correlation coefficients of the coverage values in the source distribution (inDrop), the transformed distribution (inDrop mapped by the OT model), and the target distribution (C1HT-medium).

## Notes

https://github.com/yo-aka-gene/grnet

### Competing Interest Statement

The authors have declared no competing interest.

## References

[1] F. Tang, C. Barbacioru, Y. Wang, E. Nordman, C. Lee, N. Xu, X. Wang, J. Bodeau, B. B. Tuch, A. Siddiqui, et al., “mrna-seq whole-transcriptome analysis of a single cell,” Nature methods, vol. 6, no. 5, pp. 377–382, 2009.

[2] A. Regev, S. A. Teichmann, E. S. Lander, I. Amit, C. Benoist, E. Birney, B. Bodenmiller, P. Campbell, P. Carninci, M. Clatworthy, et al., “The human cell atlas,” elife, vol. 6, p. e27041, 2017.

[3] Y. Okano, Y. Kase, and H. Okano, “A set-theoretic definition of cell types with an algebraic structure on gene regulatory networks and application in annotation of rna-seq data,” Stem cell reports, vol. 18, no. 1, pp. 113–130, 2023.

[4] A. Bookstein, V. A. Kulyukin, and T. Raita, “Generalized hamming distance,” Information Retrieval, vol. 5, pp. 353–375, 2002.

[5] P. Spirtes, C. N. Glymour, and R. Scheines, Causation, prediction, and search. MIT press, 2000.

[6] A. Ankan and A. Panda, “pgmpy: Probabilistic graphical models using python,” in Proceedings of the 14th Python in Science Conference (SCIPY 2015), Citeseer, 2015.

[7] I. Tsamardinos, L. E. Brown, and C. F. Aliferis, “The max-min hill-climbing bayesian network structure learning algorithm,” Machine learning, vol. 65, pp. 31–78, 2006.

[8] J. Zhang, J. Jiao, et al., “Molecular biomarkers for embryonic and adult neural stem cell and neurogenesis,” BioMed research international, vol. 2015, 2015.

[9] L. McInnes, J. Healy, N. Saul, and L. Großberger, “Umap: Uniform manifold approximation and projection,” Journal of Open Source Software, vol. 3, no. 29, p. 861, 2018.

[10] M. D. Luecken and F. J. Theis, “Current best practices in single-cell rna-seq analysis: a tutorial,” Molecular systems biology, vol. 15, no. 6, p. e8746, 2019.

[11] S. M. Lundberg and S.-I. Lee, “A unified approach to interpreting model predictions,” in Advances in Neural Information Processing Systems 30 (I. Guyon, U. V. Luxburg, S. Bengio, H. Wallach, R. Fergus, S. Vishwanathan, and R. Garnett, eds.), pp. 4765–4774, Curran Associates, Inc., 2017.

[12] P. Qiu, “Embracing the dropouts in single-cell rna-seq analysis,” Nature communications, vol. 11, no. 1, p. 1169, 2020.

[13] L. Zappia, B. Phipson, and A. Oshlack, “Splatter: simulation of single-cell rna sequencing data,” Genome biology, vol. 18, no. 1, p. 174, 2017.

[14] F. Puhm, A. Laroche, and E. Boilard, “Diversity of megakaryocytes,” Arteriosclerosis, Thrombosis, and Vascular Biology, vol. 43, no. 11, pp. 2088–2098, 2023.

[15] T. H. Kim, X. Zhou, and M. Chen, “Demystifying “drop-outs” in single-cell umi data,” Genome biology, vol. 21, no. 1, p. 196, 2020.

[16] K. Choi, Y. Chen, D. A. Skelly, and G. A. Churchill, “Bayesian model selection reveals biological origins of zero inflation in single-cell transcriptomics,” Genome biology, vol. 21, pp. 1–16, 2020.

[17] E. Mereu, A. Lafzi, C. Moutinho, C. Ziegenhain, D. J. McCarthy, A. Álvarez-Varela, E. Batlle, n. Sagar, D. Gruen, J. K. Lau, et al., “Benchmarking single-cell rna-sequencing protocols for cell atlas projects,” Nature biotechnology, vol. 38, no. 6, pp. 747–755, 2020.

[18] C. Wu, A. Mark, and A. I. Su, “Mygene. info: gene annotation query as a service,” bioRxiv, p. 009332, 2014.

[19] C. R. Harris, K. J. Millman, S. J. Van Der Walt, R. Gommers, P. Virtanen, D. Cournapeau, E. Wieser, J. Taylor, S. Berg, N. J. Smith, et al., “Array programming with numpy,” Nature, vol. 585, no. 7825, pp. 357–362, 2020.

[20] W. McKinney et al., “Data structures for statistical computing in python,” in Proceedings of the 9th Python in Science Conference, vol. 445, pp. 51–56, Austin, TX, 2010.

[21] F. A. Wolf, P. Angerer, and F. J. Theis, “Scanpy: large-scale single-cell gene expression data analysis,” Genome biology, vol. 19, pp. 1–5, 2018.

[22] R. Vink, “Polars: Blazingly fast dataframes in rust, python, node.js, r, and sql.” https://github.com/pola-rs/polars, 2024. Version 0.20.10.

[23] J. D. Hunter, “Matplotlib: A 2d graphics environment,” Computing in Science & Engineering, vol. 9, no. 3, pp. 90–95, 2007.

[24] M. L. Waskom, “seaborn: statistical data visualization,” Journal of Open Source Software, vol. 6, no. 60, p. 3021, 2021.

[25] G. Ke, Q. Meng, T. Finley, T. Wang, W. Chen, W. Ma, Q. Ye, and T.-Y. Liu, “Lightgbm: A highly effcient gradient boosting decision tree,” Advances in neural information processing systems, vol. 30, 2017.

[26] T. Akiba, S. Sano, T. Yanase, T. Ohta, and M. Koyama, “Optuna: A next-generation hyperparameter optimization framework,” in Proceedings of the 25th ACM SIGKDD international conference on knowledge discovery & data mining, pp. 2623–2631, 2019.

[27] F. Pedregosa, G. Varoquaux, A. Gramfort, V. Michel, B. Thirion, O. Grisel, M. Blondel, P. Prettenhofer, R. Weiss, V. Dubourg, et al., “Scikit-learn: Machine learning in python,” the Journal of machine Learning research, vol. 12, pp. 2825–2830, 2011.

[28] S. M. Lundberg, G. Erion, H. Chen, A. DeGrave, J. M. Prutkin, B. Nair, R. Katz, J. Himmelfarb, N. Bansal, and S.-I. Lee, “From local explanations to global understanding with explainable ai for trees,” Nature Machine Intelligence, vol. 2, no. 1, pp. 2522–5839, 2020.

[29] L. Kolberg, U. Raudvere, I. Kuzmin, J. Vilo, and H. Peterson, “gprofiler2– an r package for gene list functional enrichment analysis and namespace conversion toolset g:profiler,” F1000Research, vol. 9 (ELIXIR), no. 709, 2020. R package version 0.2.3.

[30] A. Paszke, S. Gross, S. Chintala, G. Chanan, E. Yang, Z. DeVito, Z. Lin, A. Desmaison, L. Antiga, and A. Lerer, “Automatic differentiation in pytorch,” 2017.

[31] S. Seabold and J. Perktold, “Statsmodels: econometric and statistical modeling with python.,” SciPy, vol. 7, p. 1, 2010.

[32] R. Flamary, N. Courty, A. Gramfort, M. Z. Alaya, A. Boisbunon, S. Chambon, L. Chapel, A. Corenos, K. Fatras, N. Fournier, et al., “Pot: Python optimal transport,” Journal of Machine Learning Research, vol. 22, no. 78, pp. 1–8, 2021.

[33] K. Tretyakov, “matplotlib-venn: Venn diagram plotting routines for python/matplotlib.” https://github.com/konstantint/matplotlib-venn, 2024. Version 0.11.10.

